# Plasmid conjugation drives within-patient plasmid diversity

**DOI:** 10.1101/2024.09.27.615342

**Authors:** Fan Grayson, Leo Loman, Toby Nonnenmacher, Diane Pople, Jack Pollard, David Williams, Bharat Patel, Luke Hounsome, Katie L Hopkins, Julie Robotham, Alice Ledda

## Abstract

Plasmids are well known vehicles of antimicrobial resistance (AMR) genes dissemination. Through conjugation, plasmid–encoded AMR genes are spread among neighbouring bacteria, irrespective of their strain or even their species. This process is very concerning from a public health perspective, as plasmid-borne AMR gene outbreaks are often not confined to single species or strains and are therefore more difficult to be fully uncovered. At the moment, the impact of plasmid conjugation on within-patient plasmid diversity is not well understood. In this work we will tackle the role of conjugation on within-patient plasmid diversity using a dataset of carbapenemase-producing *Enterobacterales* (CPEs). The dataset of 256 sequences from 115 patients was sampled across England over 30 months. Each patient has more than one sequence, with at least one sequence carrying an OXA-48 gene, a well-known plasmid-borne carbapenemase-encoding gene. If more than one sequence carries the OXA-48 gene, they are carried on different bacterial hosts. Using a hybrid *de novo*-on-reference assembly pipeline, we were able to reconstruct the full OXA-48 plasmid from short read sequencing data for 232 of the 256 sequences. Of the 115 patients, 83 (72%) of patients had an identical OXA-48 plasmid in two or more sequences. Only 2 patients carried very different (*>*200 SNPs) alleles of the OXA-48 plasmid, probably from separate acquisitions. Our study shows that when more than one bacterial host carrying an OXA-48 plasmid is found in a patient, it is most likely that the same plasmid has been shared via conjugation. The event of separate acquisition of different plasmids in different bacterial hosts is highly unlikely in our dataset.

**Data Statement:** We use data provided by Hopkins et al 2022 [16]. The data can be accessed from the National Center for Biotechnology Information (NCBI) and can be found at Bioproject Accession no. PRJNA788733. None of the data used was synthetically generated.

**Impact Statement:** Conjugative plasmids are well known vessels of horizontal gene transfer, with a prominent role in the spread of antimicrobial resistance genes among different bacterial species or strains. At the epidemiological level, conjugation combined with sequencing a single colony per patient, results in plasmids outbreaks carrying antimicrobial resistance genes being found in different bacterial species and strains in different patients, potentially eluding surveillance protocols based on same bacterium/same resistance scheme. In this study we analyse within-patient plasmid diversity in a dataset with more than one sequence per patient. Only two patients show clear genomic signs of separate plasmids acquisition, while 83 patients share identical plasmids in different bacterial hosts. This points out to a very strong role of plasmid conjugation in shaping within-patient plasmid diversity.

## 1 Introduction

From a public health perspective, plasmid conjugation is a very concerning process. Plasmid conjugation is the process by which plasmids replicate themselves and spread to neighbouring bacteria [50], facilitating the sharing of genetic material between unrelated bacterial cells [14; 40; 15]. Conjugation is one of the processes widely known as horizontal gene transfer (HGT) and can occur across bacterial strains, species, or genera [1; 2; 9; 21]. Notably, plasmid conjugation enables the dissemination of plasmid-encoded resistance genes among various bacterial species within the same patient [11; 32; 29; 51].

This poses a significant challenge in investigating plasmid-mediated Antimicrobial Resistance (AMR) outbreaks. Typically, AMR outbreaks are characterised by an excess of a specific resistance above a predefined “threshold” value in a specific bacterial host. However, when resistance is plasmid-encoded, it can easily be found in multiple bacterial hosts in the same patient. Consequently, sampling a single colony per patient may result in the resistance appearing in different strains in different patients, solely due to the conjugation ability of the plasmid, not as a pre-existing property of the pathogen. In such cases, AMR outbreak characterisation based on the single host-single resistance approach may only count sequences where the resistance is found in the specific outbreak bacterium, potentially misclassifying other sequences as non-outbreak-related.

Phylogenetics is a useful tool in tracking outbreaks of diseases, and involves analysing the genetic relatedness of outbreak isolates based on their predicted evolutionary relatedness. With epidemiological data, this can provide a better understanding of how outbreaks develop [47], and in conjunction with comparative genomics, can even help to understand underlying molecular mechanisms and vulnerabilities of the pathogen [47; 2].

In the context of plasmid-encoded AMR outbreaks, standard, substitution based phylogenetic analysis of bacterial hosts proves ineffective, as the presence of the plasmid represents an incidental occurrence, rather than the primary evolutionary driver of strains’ divergence. Conducting phylogenetic analysis on the plasmid itself, we must consider that plasmids are very ‘short’ in length, and, even considering conjugation, do not replicate often enough during a hospital outbreak to get enough neutral mutations to resolve a full tree [26]. However, plasmids can still accumulate enough adaptive mutations to resolve a full tree to learn about the plasmid’s evolutionary history beyond the simple plasmid typing and resistance identification strategy [26]. Such a tree cannot be used to establish when the plasmid clone originated, since it does not contain time related information. However, it can still be used to identify groups of related sequences (*’subclades’*), which, in connection with epidemiological features, such as location and bacterial host, specific to each subclade, can still provide a better understanding of outbreak dynamics.

Currently, the main problem with widespread usage of plasmid phylogenetics is plasmid reconstruction [10]. Short read sequencing is not ideal for reconstructing plasmid sequences, as they are circular, often incorporate repeated sequences longer than the read length thus are difficult to resolve, and it is not easy to account for all events of gene acquisition and loss in the subsequent analysis. Until recently, long read sequencing technology has been insufficiently accurate for useful phylogenetic analysis, although this can be solved using hybrid sequencing. While technology is rapidly improving, in the case of a very well-conserved plasmid such as pOXA-48, we can still attempt plasmid reconstruction from short read sequencing data [26].

Carbapenem resistance genes (e.g. OXA-48-like, KPC, NDM, and VIM families) are often encoded on plasmids [47; 26; 37]. Carbapenems are ‘antibiotics of last resort’ against multidrug-resistant infections [4]. In recent years, carbapenem-resistant infections have become more common, prompting WHO (World Health Organization) to classify carbapenem resistant Gram-negatives as a critical priority for new antibiotic development [33]. Gaining a better understanding of how plasmids carrying carbapenem resistance genes spread is therefore of paramount importance to devise effective strategies to curb their spread.

While plasmid-encoded AMR genetic determinants are spread by plasmid conjugation into different bacterial hosts, the rate at which conjugation happens at the epidemiological level is still quite unclear. Many experiments have been done to determine which plasmids conjugate more often than others, and how often they conjugate, but their results are difficult to extrapolate to the epidemiological context [18; 22].

Knowing how often we expect to find the same plasmid-encoded AMR genetic determinant in a different bacterial host in the same patient can be useful to fine-tune public health policies for plasmid-encoded AMR surveillance protocols and outbreak investigation. In particular, assessing whether the practice of sequencing a single sequence per patient can be too restrictive, and lead to an underestimation of the outbreak size. For this reason, we analysed a dataset of *Enterobacterales* sequences, chosen to have at least two sequences per patient with at least one carrying an OXA-48 resistance gene. We used this dataset to shed light on how often finding the same plasmid-encoded resistance in a patient is due to within-patient conjugation, and how often it is due to a separate acquisition of a bacterium carrying the same resistance gene, or even a different one. We reconstructed the pOXA-48 plasmid sequences present in each sequence. By studying the differences in its genetic sequence, we can tell if the plasmid is the same (hence resulting from conjugation) or different (hence resulting from separate acquisition). We use this classification to estimate the extent of conjugation and how often we can expect to find in a same patient plasmids variants so distant that can be explained by separate acquisitions.

## 2 Methods

### 2.1 Overview

We studied a dataset of within-patient bacterial sequences carrying the OXA-48 gene to better understand the role of conjugation in within-host plasmid diversity. We had an underlying hypothesis: if we find an identical plasmid in different bacterial hosts in a single patient, then the most probable scenario is that the plasmid has been exchanged between the two bacterial hosts through conjugation. If, on the other hand, we find a different variant of the plasmid in two different bacterial hosts, the patient has probably acquired two different bacterial hosts each independently carrying a variant of the plasmid. We built a pipeline to get a reliable plasmid sequence from short read sequencing. The pipeline was developed to minimize the chance of missing small insertions and deletions during assembly. We then aligned the plasmid sequences to a reference sequence (JN626286 [36]) and obtained a tree (rooted on the reference plasmid). We used the tree to define the different plasmid variants and studied the epidemiological data in the context of these variants and, more generally, the genetic data. We characterised the dataset’s mutation pattern with a focus on the within-patient versus the between-patient pattern. Finally, we used statistical modelling to identify the differences in the distributions of mutations in the within-patient and between-patient pairs of sequences. A full description of the methods is given in supplementary methods. All the scripts used to analyse data and run programs (on a high-performance computing cluster) are available on https://github.com/ukhsa-collaboration/within-patient-pOXA48-conjugation

### 2.2 Dataset selection, epidemiological data collection and sequencing

Sequence selection and criterion for sequencing is described in full elsewhere [16]. To summarise the selection and criterion, the sequences in this study were derived from bacterial isolates submitted voluntarily by diagnostic laboratories in England to UKHSA’s (formerly Public Health England, PHE) Antimicrobial Resistance and Healthcare Associated Infections (AMRHAI) Reference Unit between 1 January 2014 and 30 June 2016, for investigation of carbapenem resistance. The first referred bacterial isolate for each patient was sequenced if positive for one or more acquired carbapenemase genes. Subsequent bacterial isolates per patient were sequenced only if they carried a carbapenemase resistance gene of a different family, or if they carried the same resistance gene but in a different bacterial host. We selected all the genomes from patients who had at least one OXA-48 positive bacterial isolate. For each genome we had access to the original sampling date and site, and referring laboratory; the latter were assigned to one of nine geographical areas based on the laboratory location [12].

### 2.3 pOXA-48 Assembly

A hybrid *de novo*-on-reference pipeline was developed to obtain a reliable sequence of the pOXA-48 plasmid. A schematic drawing of the pipeline is given in Figure 2A. The rationale for this is the following: pOXA-48 is usually a very well-conserved plasmid, but it is still subject to a small degree of variability [11]. Consensus sequence among reads mapped to a reference sequence using short sequencing reads could have missed small insertions. The *de novo* pipeline running in conjunction with the on reference pipeline provides a check for the absence of small insertions included in a contig where part of the plasmid was, as it happens in contig *r* in Figure 2B. This made the on-reference assembled plasmid more reliable. It is an approach that works very well for pOXA-48 plasmids, where only small insertions and deletions are present [11]. It is difficult to generalise this method for longer plasmids with bigger insertions and deletions, that could get assembled in a separate contig.

#### 2.3.1 The *de novo* assembly pipeline

The *de novo* pipeline first ran `SPAdes` version 3.8.0 [5] on the short reads for each sequence, then aligned the output contigs to the reference plasmid sequence (GenBank record JN626286, identical to RefSeq record NC 019154 [36]) using `mummer4` [28]. Using the contigs assembled with `plasmid SPAdes` [3] (`SPAdes` with the ‘–`plasmid`’ option) does not yield better results than without. This is possibly due to the fact that the copy number of this plasmid is not high enough to make a stark difference in sequencing coverage (that is what the ‘–`plasmid`’ option uses to better identify plasmids). The *de-novo* assembly pipeline using `SPAdes` (without the `-plasmid` option was not able to reconstruct any or enough contigs for 35 of the 256 sequences, 10 of which were able to be fully assembled by the on-reference method. Using `SPAdes` without the ‘–`plasmid`’ option produced contigs for 218 out of the 232 plasmids that were able to be fully assembled by the on-reference method.

#### 2.3.2 The on-reference assembly pipeline

The on-reference assembly pipeline consisted of mapping short reads to the reference sequence with `Bowtie2` version 2.2.5 [24; 25], then calling the consensus sequence of the mapped reads relative to the reference using `samtools–consensus` [8].

#### 2.3.3 Comparison between the two pipelines

The Venn diagram in Figure 3 shows the comparison between the results of the two pipelines. The on-reference assembly pipeline was able to fully assemble 232 plasmids out of our 256 sequences. The *de-novo* assembly pipeline using `plasmid SPAdes` was not able to reconstruct any or enough contigs for 30 of the 256 sequences - 10 of these 30 plasmid sequences were able to be fully assembled by the on-reference method.

As a comparison we run Plasmid Finder. All the *de novo* contigs files obtained from `SPAdes` were run through `PlasmidFinder` [7]. Resulting `.json` files were parsed using `R` [39]. It found an IncL plasmid in 221 of the 256 sequences.

### 2.4 Bacterial Host Identification

Bacterial host identification protocol has been described elsewhere [16] and confirmed using the program `MLST` [42; 19] on whole genome contigs obtained from `SPAdes`.

### 2.5 Alignment and phylogenetic reconstruction

The on-reference assembled sequences were gathered in a single file together with the reference genome and aligned using `MAFFT` [20].

We developed an algorithm to mask hypervariable regions (see supplementary methods for details). Briefly, for every position in the alignment, if gaps were found in more than 10% of the sequences at that position, we masked all the mutations in all the sequences in the 50 bp region adjacent to the gap. If gaps were found in less than 10% of the sequences at that position, we masked the mutations in the 50 bp region adjacent to the gap only in the sequences affected by the gap. The rationale for this masking strategy is that if we find a gap in more than 10% of the sequences, we assume that we are dealing with a variable region that may be poorly aligned. However, if we find gaps in less than 10% of the sequences, we assume that the problem is in the genome sequence quality but not in the region.

This masking strategy yielded 247 mutations to resolve the tree.

The masked alignment was used to reconstruct the phylogenetic tree with IQtree version 1.6.12-Linux [31] under the GTR substitution model [43] with the discrete Gamma model [52] with the default 4 rate categories to correct for rate heterogeneity across sites. The reference plasmid (JN626286 [36]) was used to root the tree as it is the first pOXA48 plasmid isolated in 2001, so an ancestor to our dataset.

Figures for all trees were produced using the following `R` [39] packages: `ape` [35], `ggtree` [53], `treeio` [48], and `ggnewscale` [6].

### 2.6 Within-patient diversity

Within-patient diversity was computed using a script in `R` [39] (see supplementary methods and github repository for detail).

### 2.7 Plasmid ancestral state reconstruction

The epidemic clade improved alignment and tree were fed to the program `augur ancestral` [17], that uses maximum likelihood to return the ancestral sequence at each node. Results were analysed and plotted using `R` [39] (see supplementary methods and github repository).

### 2.8 Comparing within-patient and between-patient diversity

To analyse the distribution of sequences from a same patient in the tree, specifically, the likelihood that each clade contains a pair of sequences from the same patient, statistical tests were performed on the genetic distance between pairs of different plasmid sequences. The pairwise distances are split into two groups:

**Population 1:** All distances between pairs of different plasmid sequences excluding the ones in Population 2
**Population 2:** Distances of different plasmid sequence pairs where both sequences in the pair come from the same patient

The hypothesis test is as follows:

**H**_0_: Population 1 and 2 have the same distribution of genetic distance
**H**_1_: Population 1 and 2 have different distributions of genetic distance We performed two non-parametric hypothesis tests, which were the

Mann-Whitney test [27; 30] and the Wilcoxon Signed-Rank test [49]. These tests were performed in python utilising the `scipy` package version 1.10.1 [46] (using the `stats.mannwhitneyu` and `stats.median test` functions). The results of these are in Table 2a.

A parametric hypothesis test was undertaken by using a Negative Binomial model [34]. This was performed in `R` using the `mass` package version 7.3.57 [45]. We chose this model based on the general distribution of the pairwise distances (heavily skewed left) and a few assumptions (genetic distance has to be a non-negative integer value – this is count data, and we cannot have fewer than 0 SNPs, and there is theoretically no maximum limit for genetic distance). The result of this is in Table 3a.

These same tests were run on the sequences sampled in the North West to discard the hypothesis that we saw more sequence similarity in this region because of an ongoing outbreak.

## 3 Results

### 3.1 Dataset Description

To study the within-patient variability associated with carriage of OXA-48 genes, we filtered data from a previous study [16] to only include sequences from patients that had at least two sequences with at least one identified as carrying OXA-48. The aim of this study is understanding within-host variability associated with OXA-48 carriage, and, in particular: how often we expect to find the same OXA-48 plasmid variant associated with OXA-48 carriage; how often we expect to find a different OXA-48 plasmid variant and how often we expect to find an entirely different carbapenem resistance gene.

With these requirements, the final dataset consisted of 256 sequences from 115 patients. The median number of sequences per patient was 2 (mean 2.22). The distribution of the number of sequences per patient is shown in Figure 1A. The geographical distribution of the referring laboratory location is given in Figure 1B and the sampling dates are shown in Figure 1D.

**Figure 1:**
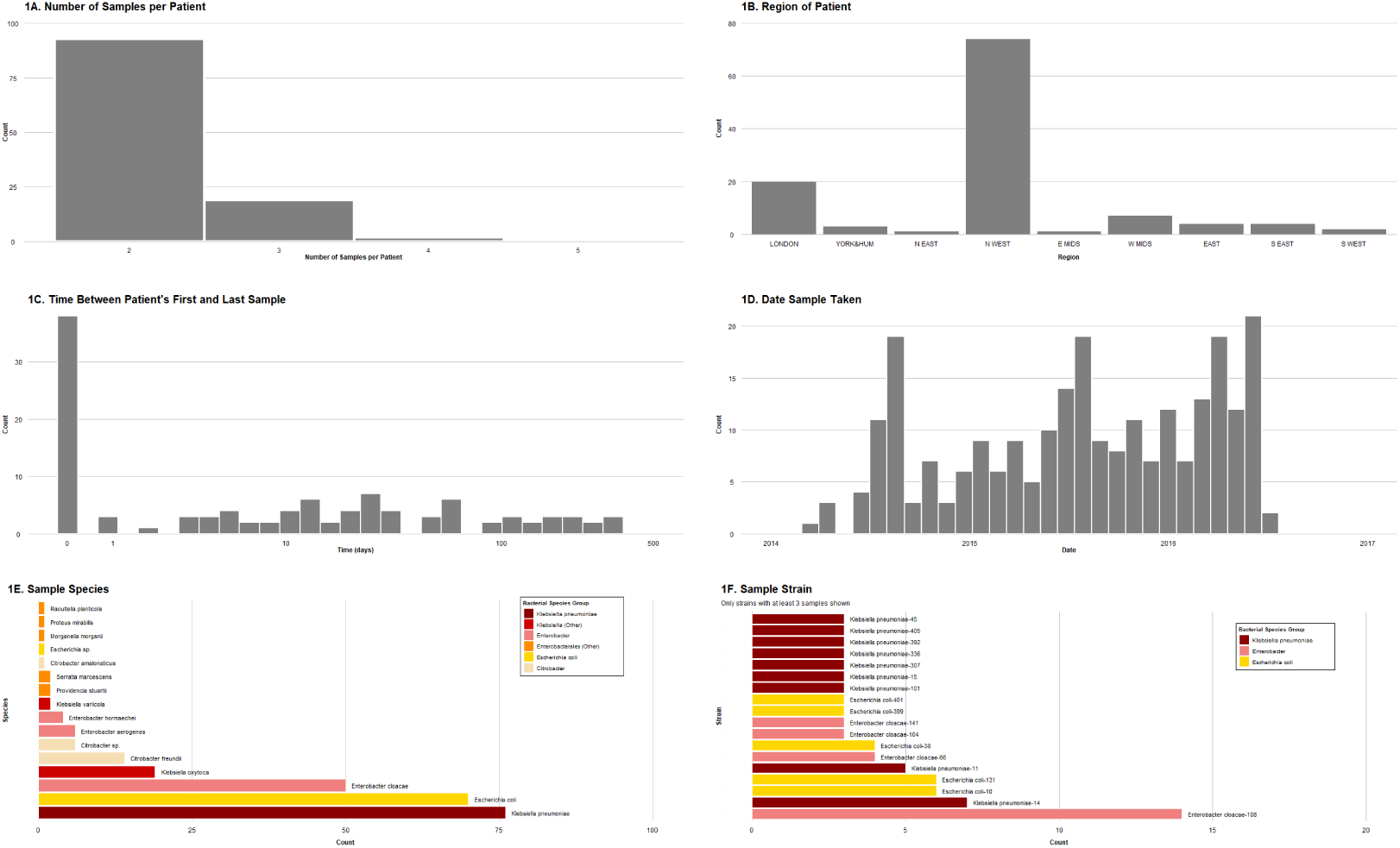
Statistics of the dataset. **A.** Number of sequences per patient. **B.** Geo-graphical region of origin of the sequence. **C.** Within-patient time interval between sequences. **D.** Date of sampling of each sequence. **E.** sequence species (coloured by species) **F.** sequence strain (coloured by species).

Sequences originating from the same patient were taken at a variable range of time intervals, ranging from the same day up to a one year interval (Figure 1C). A third (38/115; 33.0%) of the patients had all their sequences taken on the same day. The median sampling interval was 9 days (mean 41 days).

Sixteen *Enterobacterales* species were represented in our dataset (Figure 1E). The most common were *Klebsiella pneumoniae* (76/256; 29.7%), *Escherichia coli* (70/256; 27.3%), *Enterobacter cloacae* (50/256; 19.5%), *Klebsiella oxytoca* (19/256; 7.4%), and *Citrobacter freundii* (14/256; 5.5%). The complete distribution of bacterial species is given in Figure 1E.

Analysis at the strain level provides a more fragmented picture of bacterial host diversity. In the case of a clonal expansion of a specific plasmid-host pair, we would expect a single bacterial strain to be highly represented. However, in our dataset no strain is highly represented among bacterial hosts, so a clonal expansion of a specific bacterial host is responsible does not explain our data. Overall, the most common strain found in the sequences is *Enterobacter cloacae* ST108 (14/256 sequences; 5.5%), followed by *Escherichia coli* ST10 (6/256; 2.3%), *Escherichia coli* ST131 (6/256; 2.3%), and *Klebsiellae pneumoniae* ST14 (6/256; 2.3%). All of them are very common strains in their species. The distribution of bacterial strain is given in Figure 1D.

Among the 70 *E. coli* sequences present in our dataset, there were 47 sequence types (ST), 36 of which were singletons. The most common *E. coli* strains were ST131 (6/70 sequences; 8.6%), a well-known epidemic and virulent clone, and ST10 (6/70; 8.6%). Other common *E. coli* strains in our dataset include ST38 (4/70; 5.7%), a strain well known to be associated with OXA-48 carriage, albeit often inserted in the chromosome [38], and ST399 (3/70; 4.3%), another strain associated with pOXA-48 carriage in the UK [26]. Seven other *E. coli* STs were represented by fewer than three sequences each.

A similar diversity of strains was found for *Klebsiella pneumoniae* species. Among the 76 *K. pneumoniae* sequences present in the dataset, there were 49 *K. pneumoniae* STs, 37 of which were singletons. The most common *K. pneumoniae* STs were ST14 (6/76; 7.9%) and ST11 (5/76; 6.6%). Both strains happen to correspond to the most abundant *Klebsiella pneumoniae* strains worldwide. Nine other *K. pneumoniae* STs were represented by two to four sequences each.

Amongst the 50 *Enterobacter cloacae* sequences present in the dataset, the most abundant ST was ST108 (14/50; 28%), and accounted for the most common ST in the entire dataset. Nineteen *E. cloacae* STs were singletons, and five other *E. cloacae* STs were represented by two to four sequences each.

### 3.2 Plasmid reconstruction from short read sequencing

The OXA-48 gene is commonly harboured in a 61881 bp (basepair) IncL/M-type plasmid [36]. While other plasmids are well known to be highly variable in their gene content, pOXA-48 is an atypical plasmid in that its gene content is largely well-conserved and evolution acts mostly in terms of point mutations or by homologous recombination [26]. Still, in large sequence datasets, some degree of variability in pOXA-48 has been observed: De la Fuente et al identified 35 pOXA-48 plasmid variants, mainly characterised by point mutations but also by deletions ranging from a few hundred bps and up to several kilobases (kb) with their consensus plasmid being 65499 bps long, *∼*3700 bps longer than the plasmid used as a reference in this paper [11].

For plasmid reconstruction, an empirical hybrid *de novo*-on-reference pipeline was developed that has been proposed elsewhere [26]. This pipeline is illustrated in Figure 2A.

**Figure 2:**
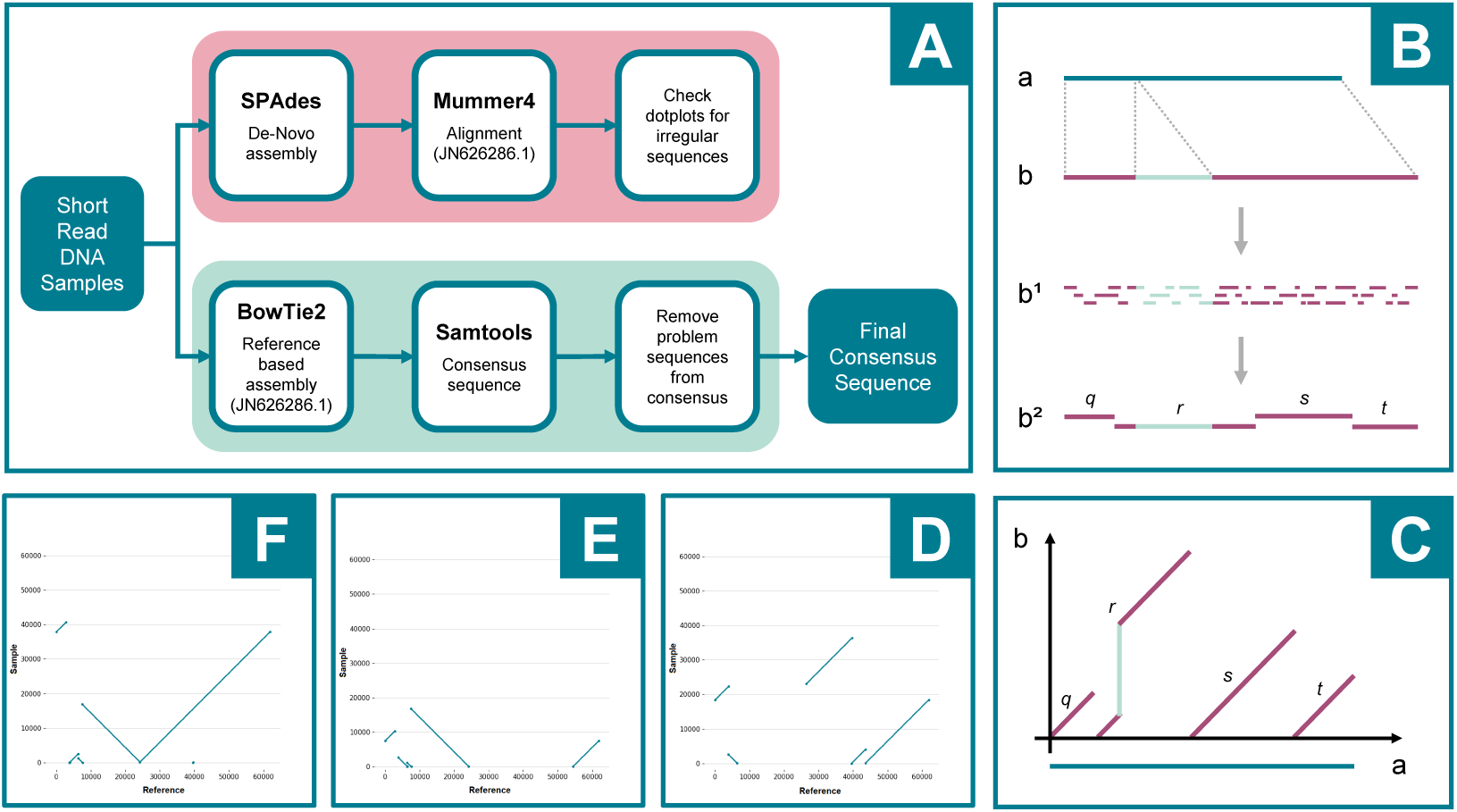
Plasmid reconstruction from short read sequencing. **A.** Flowchart describing the pipeline used (see Methods for details). **B.** Illustration of short read-based *de novo* assembly. Sequence *a* (teal) is the reference plasmid sequence obtained from NCBI (JN626286), and sequence *b* (mauve) is a putative plasmid sequence we need to reconstruct from short read sequences (*b*^1^). In this case we assume that plasmid *b* is identical to the reference sequence *a*, except for an insertion (light green). Using *de novo* based assembly, we construct contigs (*b*^2^), but we are still not able to circularise the plasmid. The contigs have been labelled *q*, *r*, *s*, and *t*. Contig *r* covers the entire insertion (light green), while contigs *q*, *s*, and *t* do not have any insertions compared to reference sequence *a*. **C.** The vignette shows how, for this case, we would be able to identify the insertion by mapping the contigs on to the reference plasmid in a dotplot. The *de novo* assembled contigs (*q*, *r*, *s*, and *t*), obtained from vignette B, are mapped to the reference sequence *a* (on the *x* axis). Contigs *q*, *s*, and *t*, each with no insertions, are mapped as unbroken diagonal lines with a slope of 1 (i.e. each base in the contig corresponds to a base in the reference and vice versa) starting at 0 on the *y* axis (contig) and at the beginning of their location in the reference on the *x* axis. Contig *r*, which contains an insertion, is also mapped diagonally starting at y=0. It is unbroken except in the area of the insertion. The insertion appears as a vertical gap (illustrated in light green), as the insertion region does not map onto the reference sequence and shifts the rest of the contig further up. **D.** Example of a dotplot for plasmid showing a deletion. **E.** Example of a dotplot for a plasmid showing a deletion and probably an inversion. **F.** Example of a dotplot for a plasmid showing a fully reconstructed plasmid (the last contig, starting from *∼*24kb, has been reconstructed in the reverse strand.

The rationale for this hybrid *de novo*-on-reference pipeline is that we expect the plasmid’s genetic content to be well-conserved. More precisely, given the specific conservation features of this plasmid, we expect almost complete synteny except for occasional small (*<*1kb) insertions. A standard reference-based reconstruction might miss insertions in the plasmid sequence (Figure 2B). To control for small insertions, we run a *de novo* assembler (Figure 2B) and then we map its contigs on the reference plasmid (Figure 2C). Considering that the whole plasmid was often covered by only a few contigs, small insertions would have been visible, mapping the contigs on the reference plasmid in the dotplots as discontinuities in the contig (*y* axis, Figure 2C), while deletions would be clearly seen on the dotplot as discontinuities on the reference (*x* axis, Figure 2D and Figure 2E). In Figure 2F, we show for comparison an example of a dotplot with no discontinuities (perfect assembly). In this case, the contig running from *∼*8kb to *∼*25kb is mapped on the reverse strand, hence the downward slope. All the dotplots were visually inspected and no sign of insertions were detected.

Of the 243 sequences in the dataset experimentally identified as carrying the OXA-48 gene, the full pOXA-48 sequence can be reconstructed for 232 sequences (95%) using this hybrid pipeline. Additionally, 6 out of the 243 sequences were experimentally identified as carrying both NDM-1 and OXA-48 genes, and 16 sequences were experimentally identified as carrying genes encoding other carbapenemase genes (see Table 1).

**Table 1:** Number of sequences with Carbapenem Resistances. (Excluding OXA-48)

| Resistance | Number of sequences |
| --- | --- |
| NDM-1 | 5 |
| KPC-2 | 5 |
| NDM-4 | 1 |
| OXA-181 | 1 |
| OXA-244 | 1 |
| VIM-4 | 1 |

By manually reviewing the dotplots, we found 3 sequences with a contig mapping on the region of the reference plasmid where the OXA-48 gene is, but no other contigs mapping on the rest of the reference plasmid. Of these 3 sequences, one (`PAT09-S020`) carried an OXA-181 gene, whose sequence differs from the OXA48 sequence by only 4 mutations, therefore mapping on the OXA-48. OXA-181 is known to be carried on a variety of non-IncL/M-type plasmids [37]. The other two sequences carried OXA-48 genes, the first from an *E. coli* ST38 (`PAT233-S505`), and the second sequence carrying OXA-48 in a *Citrobacter* (`PAT92-S204`). *E. coli* ST38 has been previously described with part of pOXA-48 inserted in its chromosome [44]. Insertion of OXA-48 in a *Citrobacter* chromosome or in a non-IncL/M-type plasmid is also described [23]. These provided a useful check that our pipeline worked properly.

As a consistency check, we run `PlasmidFinder` on the whole dataset [7] as it is the usual standard for plasmid indentification. It identified an IncL/IncM plasmid in 221 of the 256 sequences in the dataset. `PlasmidFinder` results are given in Supplementary Table.

A visual comparison between `PlasmidFinder` and each part of our hybrid pipeline findings is shown in Figure 3. `PlasmidFinder` identified a pOXA-48 plasmid in 221 out of the 256 sequences. This is compared to 226 sequences found with pOXA-48 using the *de novo* pipeline, and 232 using the reference-based pipeline. The *de novo* pipeline identified the pOXA-48 plasmid in two sequences where no other pipeline identified it. The reference-based pipeline identified the pOXA-48 plasmid in two sequences where no other pipeline identified it. `PlasmidFinder` did not exclusively identify any plasmid. Thirteen sequences were found by the *de novo* and the reference-based pipeline but not by `PlasmidFinder`. Three sequences were found by the *de novo* pipeline and `PlasmidFinder` but not by the reference-based pipeline. Two sequences were found by the reference-based pipeline and `PlasmidFinder` but not by the *de novo* pipeline. For 208 sequences out of 256, the pOXA-48 plasmid was identified using all three pipelines. All three pipelines agreed that for 18 sequences out of 256, no pOXA-48 plasmid is present.

**Figure 3:**
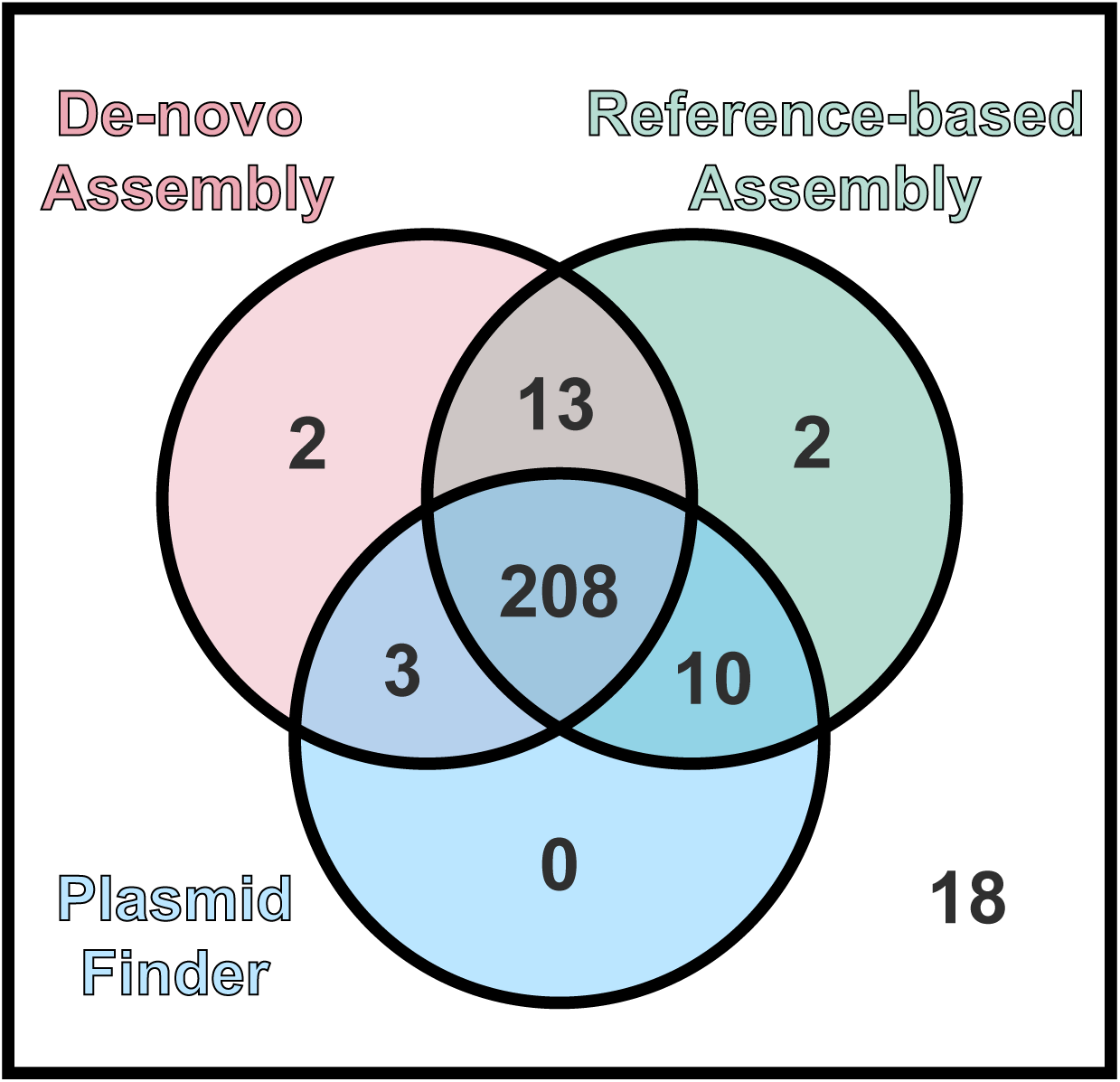
pOXA-48 Identification Venn Diagram. Venn diagram showing number of sequences where pOXA-48 is found by each of the different pipelines. The Venn diagram has been colour coded in a similar fashion to Figure 2A, with the *de novo* pipeline coloured red and the *on-reference* pipeline coloured green.

Having made sure that our pipeline was accurate enough and that no insertions were present in our dataset, all of the plasmid sequences obtained from the on-reference mapping of the reads were used to build the alignment (232/256 sequences).

### 3.3 Tree analysis

To better understand the variability of the pOXA-48 plasmid, we inferred a phylogenetic tree from all 232 of the plasmid sequences we were able to reconstruct using the reference-based pipeline, and used the reference plasmid (JN626286) as an outlier to root the tree (Figure 5).

Throughout this paper we will define a clade in a tree as sets of closely related sequences originating from a common ancestor. The tree is clearly split into clades, one containing 17 very genetically diverse sequences, and another containing 215 very closely related sequences. The former, mainly representing plasmid sequences from bacterial isolates originating in London, may be considered as a sampling of the available pOXA-48 diversity worldwide while the latter clade is clearly driven by the epidemic pOXA-48 clade expanding in some regions of the UK. From here on we will refer to the first clade as the “diverse” clade (shaded in light orange in Figure 5) and to the latter as the “epidemic” clade (shaded in light blue in Figure 5). These clades have different geographical origin: the diverse clade has mostly been sampled in London (p-value *<*0.01, Fisher exact test), while the epidemic clade has mostly been sampled in the North West of England (p-value *<*0.001, Fisher exact test). Sampling time and bacterial host are not as significantly different between the clades (bacterial host: p-value=0.012, Fisher exact test; sampling time: p-value=0.2 Kolmogorov-Smirnov test).

A closer look at the epidemic clade (Figure 6) gives a clearer view of the diversity in this clade. Within this clade, there is a cluster of 150 identical sequences, while the remaining 65 sequences in this clade display some diversity. Within this epidemic clade, there are 6 main subclades (labelled A to F), containing 4 to 13 sequences, and with sequences belonging to more than one patient. While these clades do not show clustering in terms of host species (non-significant p-value for each clade, Fisher exact test) or sampling time (non-significant p-value for each clade, Kolmogorov-Smirnov test), four of the six clades (B, D, E, and F) have a well-defined geographical origin. Clade B has a mixed geographical profile (two sequences from the South West region, one from the East of England, and one from London), but still significantly different from the overall geographic profile of the epidemic clade (p-value = 4 *×* 10*^−^*^4^, Fisher exact test), clade D is from the West Midlands (p-value *<* 10*^−^*^10^, Fisher exact test), clade E is from London (p-value = 10*^−^*^5^, Fisher exact test), and clade F has a combination of sequences from the North West and the East of England still significantly different from the whole epidemic clade (p-value = 0.0014, Fisher exact test). Clades A and C instead come from the North West, like most of the sequences in the epidemic clade, therefore the p-value from the Fisher exact test is non-significant.

Throughout the dataset, the only epidemiological feature linked to the genetic difference is the geographical origin of each sequence. This points to small outbreaks of resistance spreading in geographically limited regions, but further information (e.g., city or hospital trust) would be needed to support this hypothesis.

### 3.4 Within-patient Diversity

A footprint of within-patient conjugation is the presence of identical plasmids in different bacterial hosts within the same patient. To assess the similarity between plasmids within patients, we computed the number of single nucleotide variants found between any pair of pOXA-48 sequences from the same patient (within-patient pairwise genetic distance). In general, when comparing variability between plasmids, we call each genetic variant a “plasmid variant”. When a patient has two identical plasmids (pairwise genetic distance=0), they carry the same plasmid variant. When the patient carries two plasmids with pairwise genetic distance *>*0, we say that they carry different variants of the plasmid. Two different variants of the plasmid might be evolutionarily related to each other, meaning that they share some of the same mutations, which makes us think that one is an ancestor and the other is a descendant. On the other hand, when two different plasmid variants do not share any mutation, it is unlikely that they are evolutionarily related to each other.

Given the within-patient distribution of the sequences (see Figure 1A) we were able to compute 136 within-patient pairwise genetic distances. The distribution of these distances is given in Figure 4. As expected from the tree 5, the vast majority, 112 of 136 (82%) of the sampled patients yielded identical plasmid sequences (genetic distance = 0). Of the 136 total sampled patients, 13 (10%) plasmid sequences differ by a handful of mutations (1-6 SNPs). A further five pairwise genetic distances are in the tens of SNPs and a further 3 distances are in the hundreds of SNPs. The average within-patient pairwise genetic distance is 9.2 SNPs, the median is 0 SNPs.

**Figure 4:**
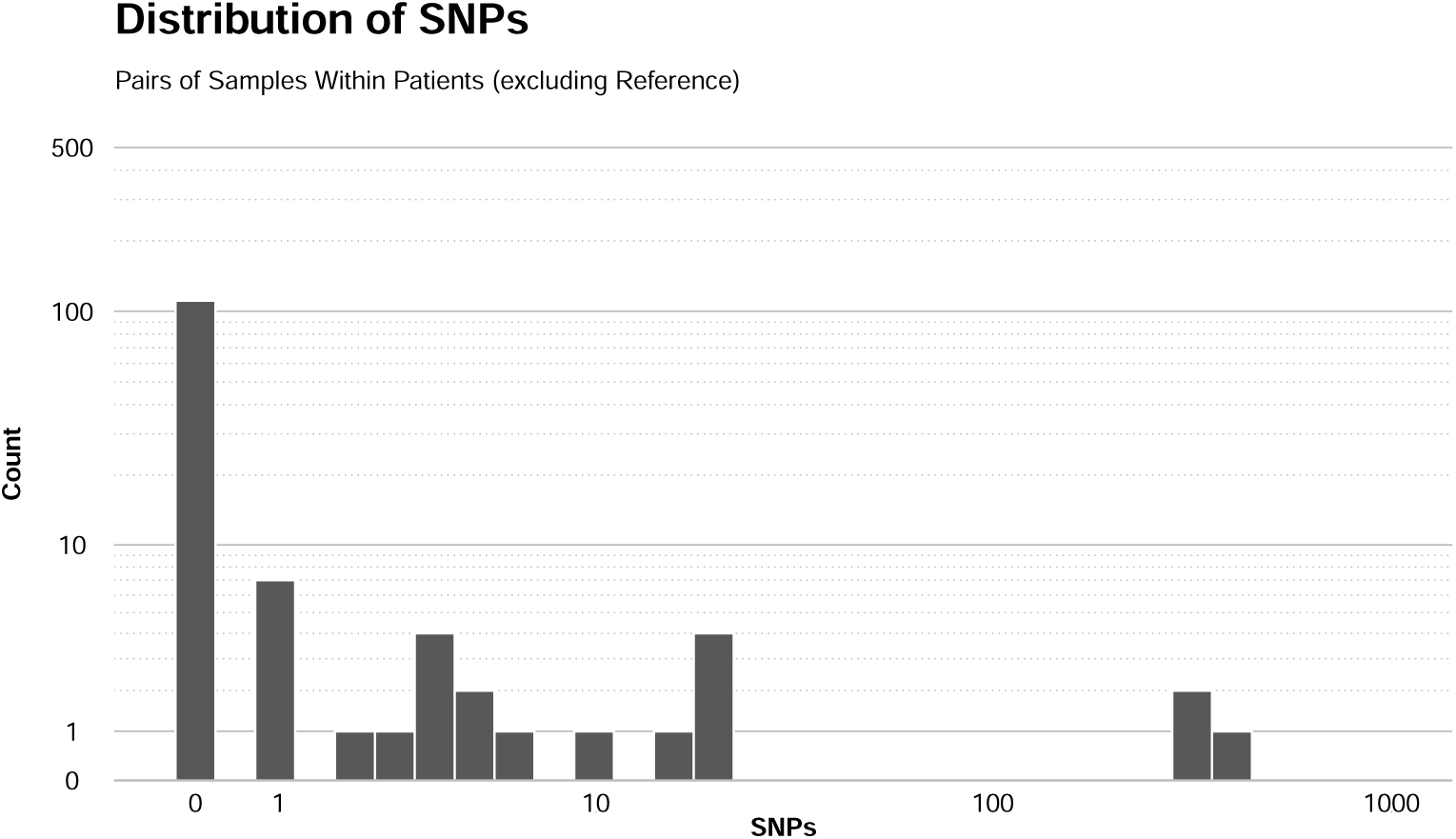
Histogram of within-patient genomic distances.

**Figure 5:**
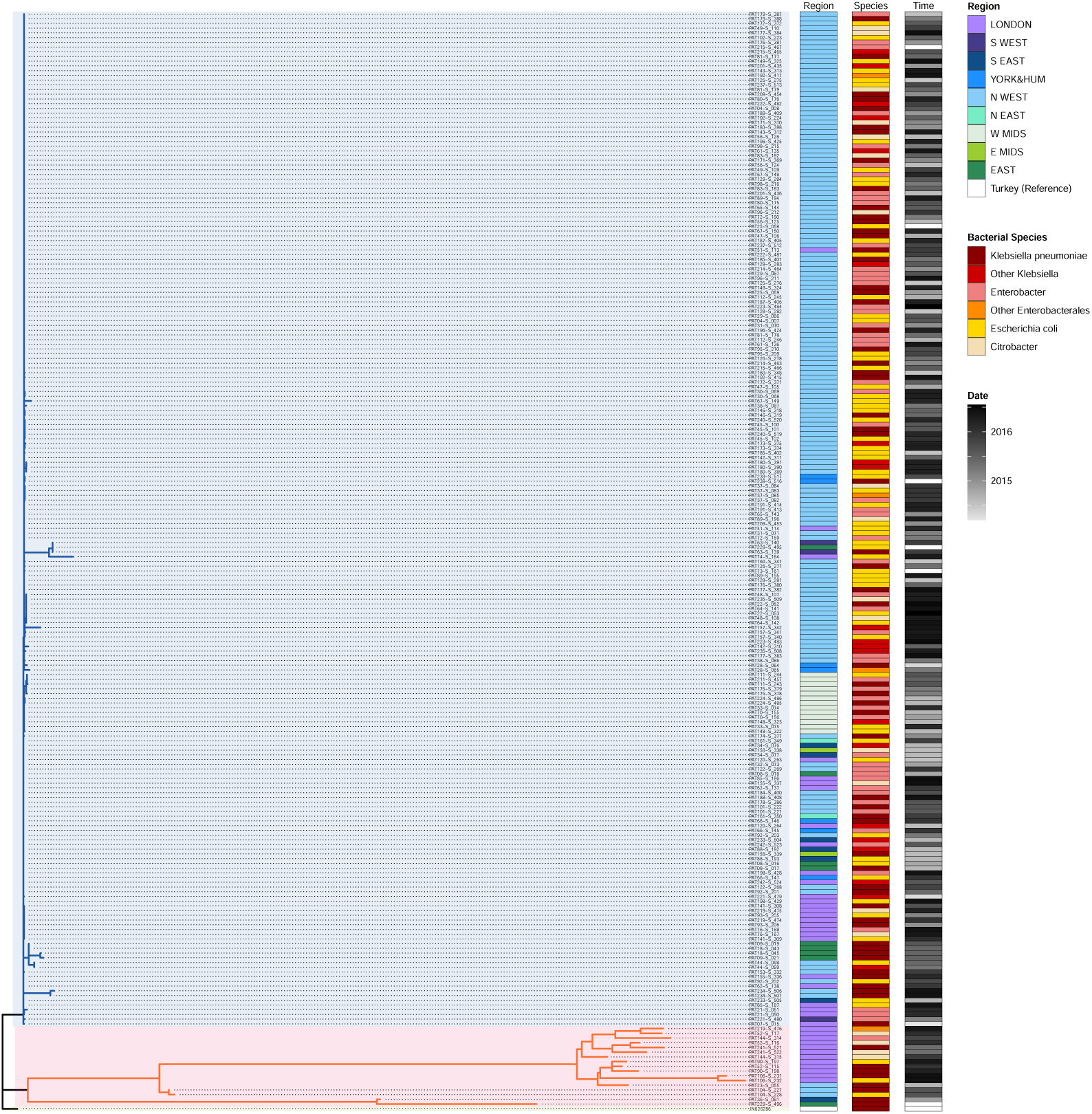
Phylogenetic Tree. All pOXA-48 sequences. Phylogenetic Tree and Analysis of the pOXA-48 sequences. The phylogenetic tree is rooted at the reference plasmid JN626286. It is split at its base into two main clades, shaded in light orange (the “diverse” clade) and light blue (the “epidemic” clade) for easier identification. The three colour-coded columns to the right of the tree show the epidemiological data for each sequence. The first column labelled “Region” (shades of blue, purple, and green) indicates the region where the sequence was collected. The second column labelled “Species” (shades of orange, red, and yellow) indicates the bacterial host species. The third column labelled “Time” (shades of grey) indicates the sampling time.

**Figure 6:**
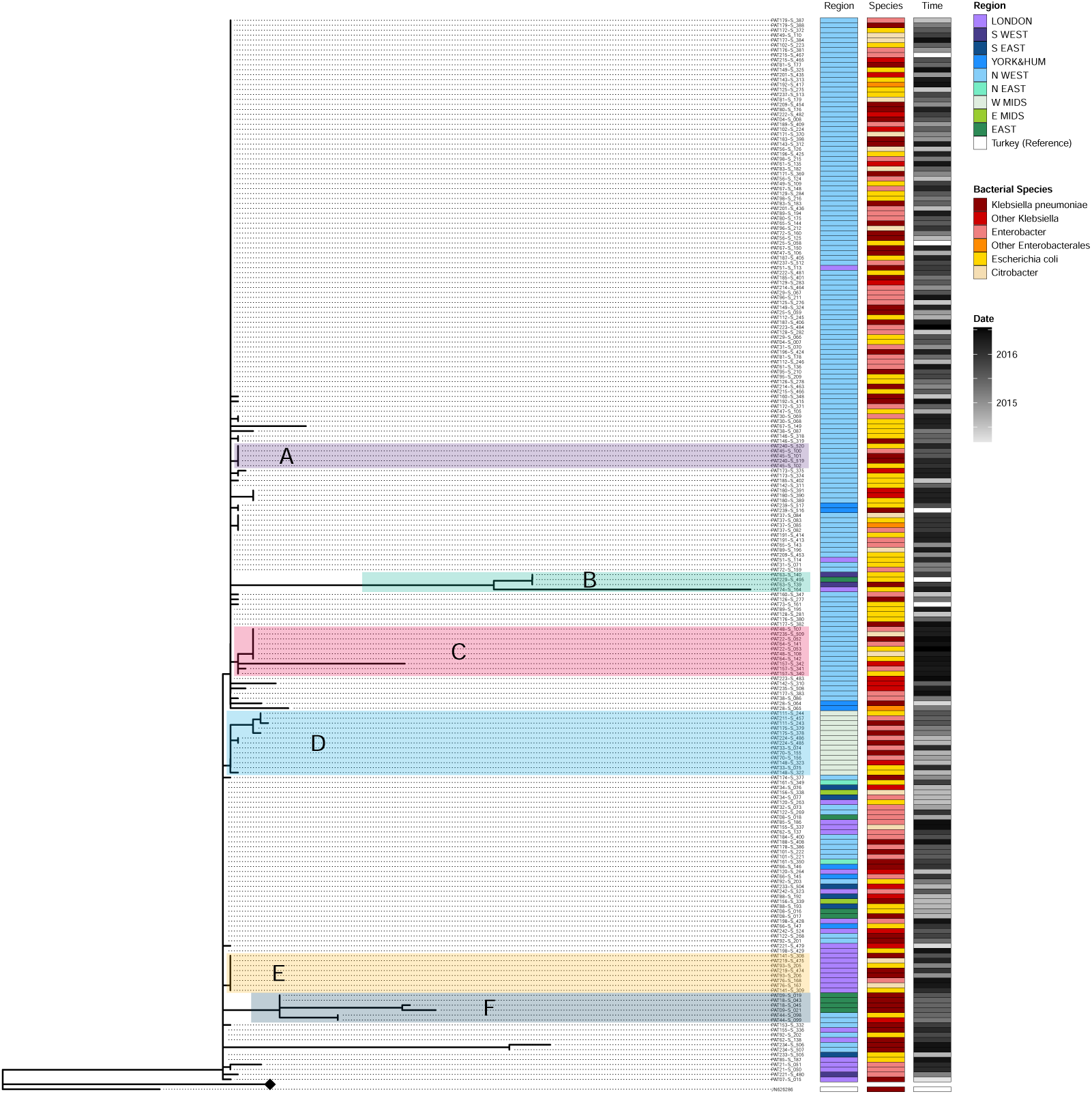
Phylogenetic Tree. Subclades within Epidemic Clade. Detail of the epidemic clade in the tree (a portion of the tree has been collapsed and represented with a diamond node). Columns have the same color coding as in Figure 5. The most populated clades are highlighted in different coloured shades and labelled A-F.

When looking at the within-patient diversity, we find the same OXA-48 plasmid variant in two or more sequences in 85 of the 115 patients in our dataset (74%). In 83 of these 85 patients (72% of the total) all sequences contained the same variant indicating no pOXA-48 diversity. In two patients, `PAT157` and `PAT219`, from which three OXA-48-positive bacterial isolates were each isolated, we find two identical variants and one different variant pOXA-48 plasmid. Eighteen of the 115 (16%) patients carry at least two different plasmid variants. There is no significant difference in sampling time interval in the carriage of identical plasmid alleles or different plasmid variants (Kolmogorov-Smirnov test).

Only two patients have sequences from both the epidemic and the ‘diverse’ clade (`PAT229` and `PAT219`, seen in Figure 7). Since the two clades are so distinct, this can only happen if the plasmids have been separately acquired. In particular, `PAT219` is a textbook example of separate acquisition, as both the different alleles we find in each sequence have another identical plasmid found elsewhere on the tree in sequences from other patients and different bacterial hosts.

**Figure 7:**
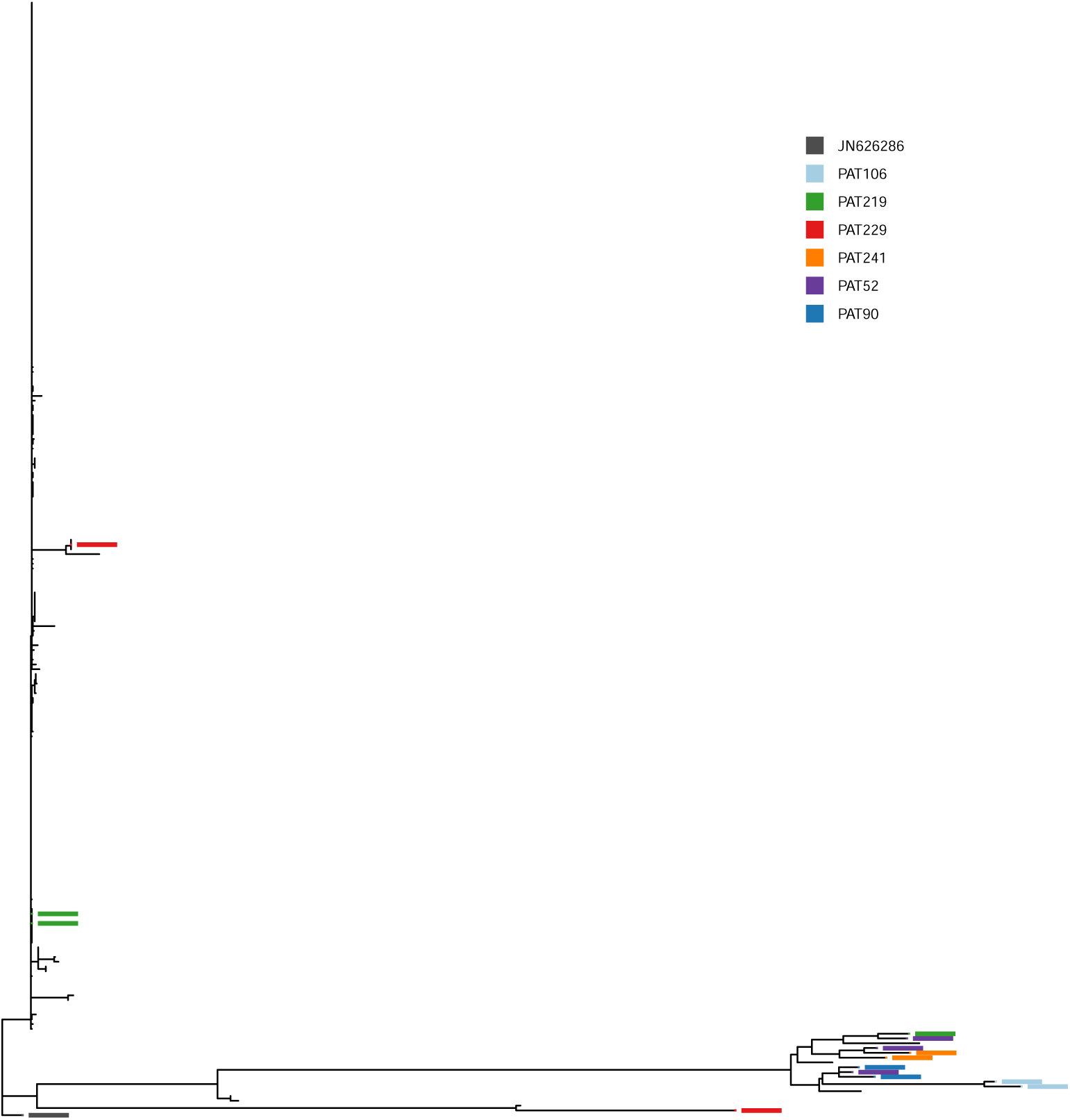
Phylogenetic Tree. Patients with sequences containing distinct pOXA-48 alleles and at least one sequence in ‘diverse’ clade. Patients with sequences containing distinct alleles of the pOXA-48 plasmid, and with at least one sequence in the non-epidemic *diverse* clade. These are good candidates for having acquired the different plasmids on different occasions. Node highlight colours correspond to each patient.

The rest of the sequences show some diversity, and when plotted on the tree, do not show clear footprints of separate acquisitions. In the case of within-patient evolution, we expect one of the sequences to sit at the base of the clade and the other at the tip. Examples of such dynamics are `PAT157`, `PAT09`, and `PAT18` in Figure 8. For other sequences, like those from `PAT106`, `PAT241`, and `PAT90` in Figure 7, it is clear that the sequences are somehow related through their evolution, but more data would required to determine whether this is due to within-patient evolution or due to a wide transmission bottleneck from an evolving source.

**Figure 8:**
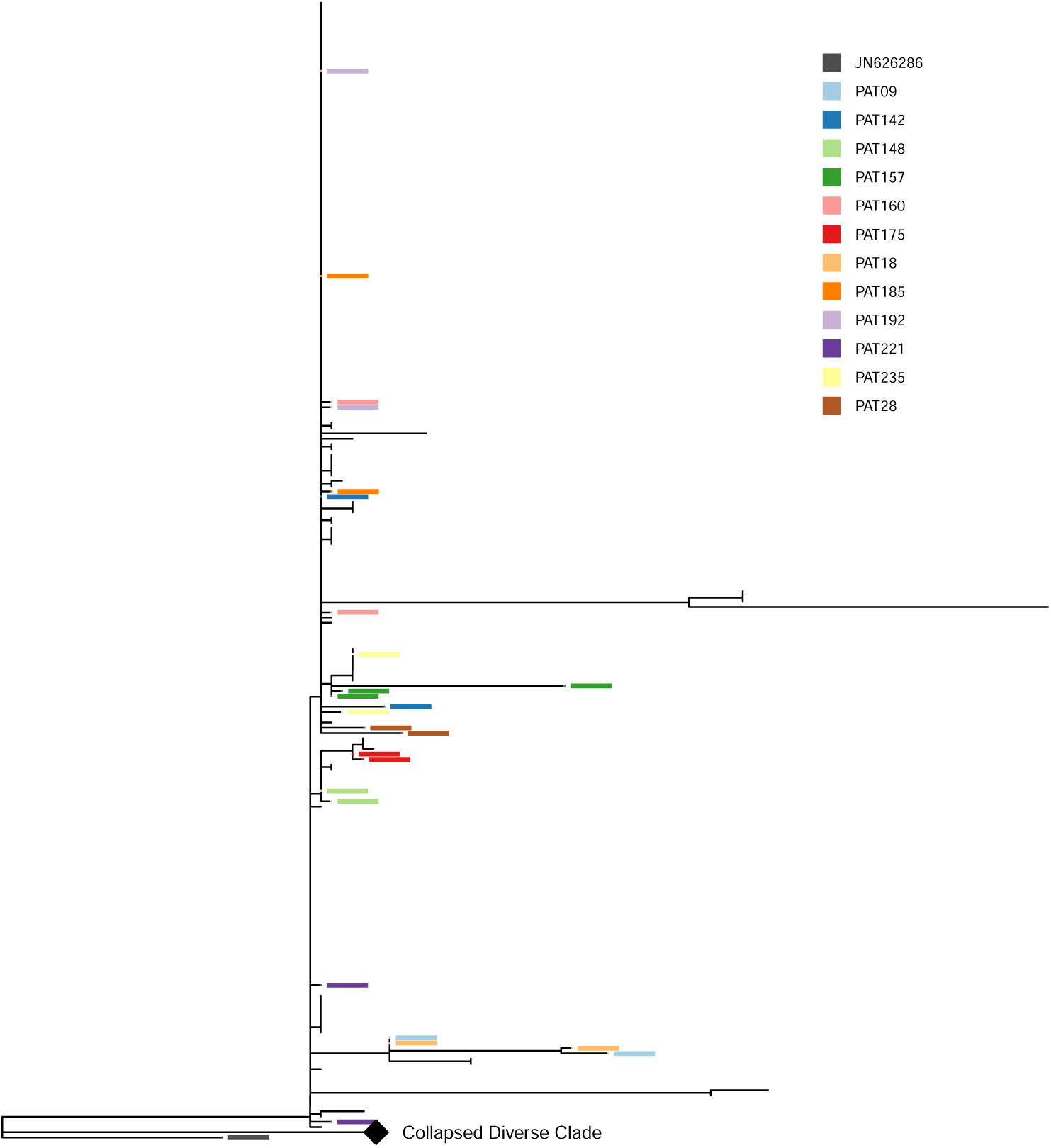
Phylogenetic Tree. Patients with sequences containing distinct pOXA-48 alleles and no sequences in ‘diverse’ clade. Patients with sequences containing distinct alleles of the pOXA-48 plasmid, and with *no sequences* in the non-epidemic *diverse* clade. These are good candidates for having acquired the different plasmid in a different occasion. Node highlight colours correspond to each patient. The non-epidemic ‘diverse’ clade has been collapsed and is represented with a diamond node.

### 3.5 Comparing Within-patient and Between-patient Diversity

If the plasmids found within the same patient have been acquired separately from a common set of plasmids (in this case our full dataset), we would expect this to reflect in the distribution of their length. Namely, we would expect the distribution of within-patient plasmid lengths to be non-statistically significantly different from the distribution of plasmid length in the entire dataset. If, on the other hand, conjugation has a role in plasmid spread, plasmids found within-patient would not be randomly sampled from the entire dataset. We would therefore expect the within-patient distribution of plasmid lengths to be statistically significantly different from the global distribution of plasmid lengths in the dataset. The results of the tests can be found in Table 2b. These tests all show that there is a statistically significant difference between the within-patient plasmid subset and the whole dataset. The tests all indicate that we can reject the null hypothesis which is that the distributions of the two sets are the same, suggesting that conjugation does have a role in plasmid spread.

**Table 2:**
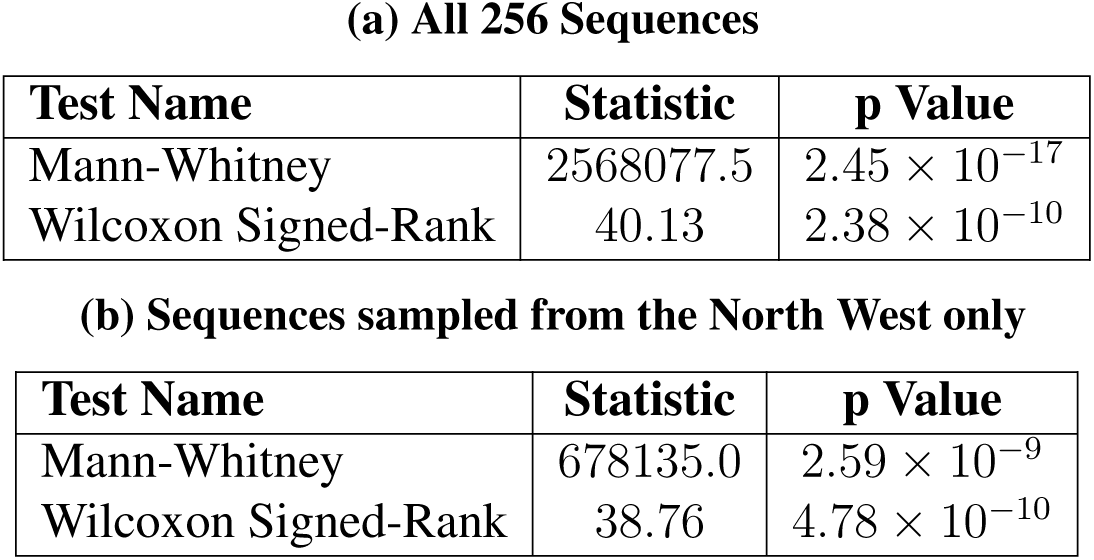
Results of Non-Parametric Hypothesis Tests. Results of Pairwise Genetic Distance Non-Parametric Hypothesis Tests (see Section 3.5).

| Test Name | Statistic | p Value |
| --- | --- | --- |
| Mann-Whitney | 2568077.5 | $2.45 \times 10^{-17}$ |
| Wilcoxon Signed-Rank | 40.13 | $2.38 \times 10^{-10}$ |

**(b) Sequences sampled from the North West only**
| Test Name | Statistic | p Value |
| --- | --- | --- |
| Mann-Whitney | 678135.0 | $2.59 \times 10^{-9}$ |
| Wilcoxon Signed-Rank | 38.76 | $4.78 \times 10^{-10}$ |

To exclude that the increased similarity in within host plasmids in the epidemic clade was due to sampling from an ongoing outbreak in that region, the same tests have been performed on sequences sampled from the North West only. In an ongoing outbreak we would expect a recent clonal expansion of the outbreak plasmid clone. Being recent, they would not have had time to acquire mutations (random or adaptive alike), therefore we expect outbreak sequences to be more similar than non-outbreak sequences. P-values were still significant, meaning that, despite being sampled in an outbreak, the within-patient similarity was different from the between-patient similarity (see table 3b). This result points again to the role of plasmid conjugation in within-patient similarity (as opposed to random sampling from the pool of plasmids available).

**Table 3:**
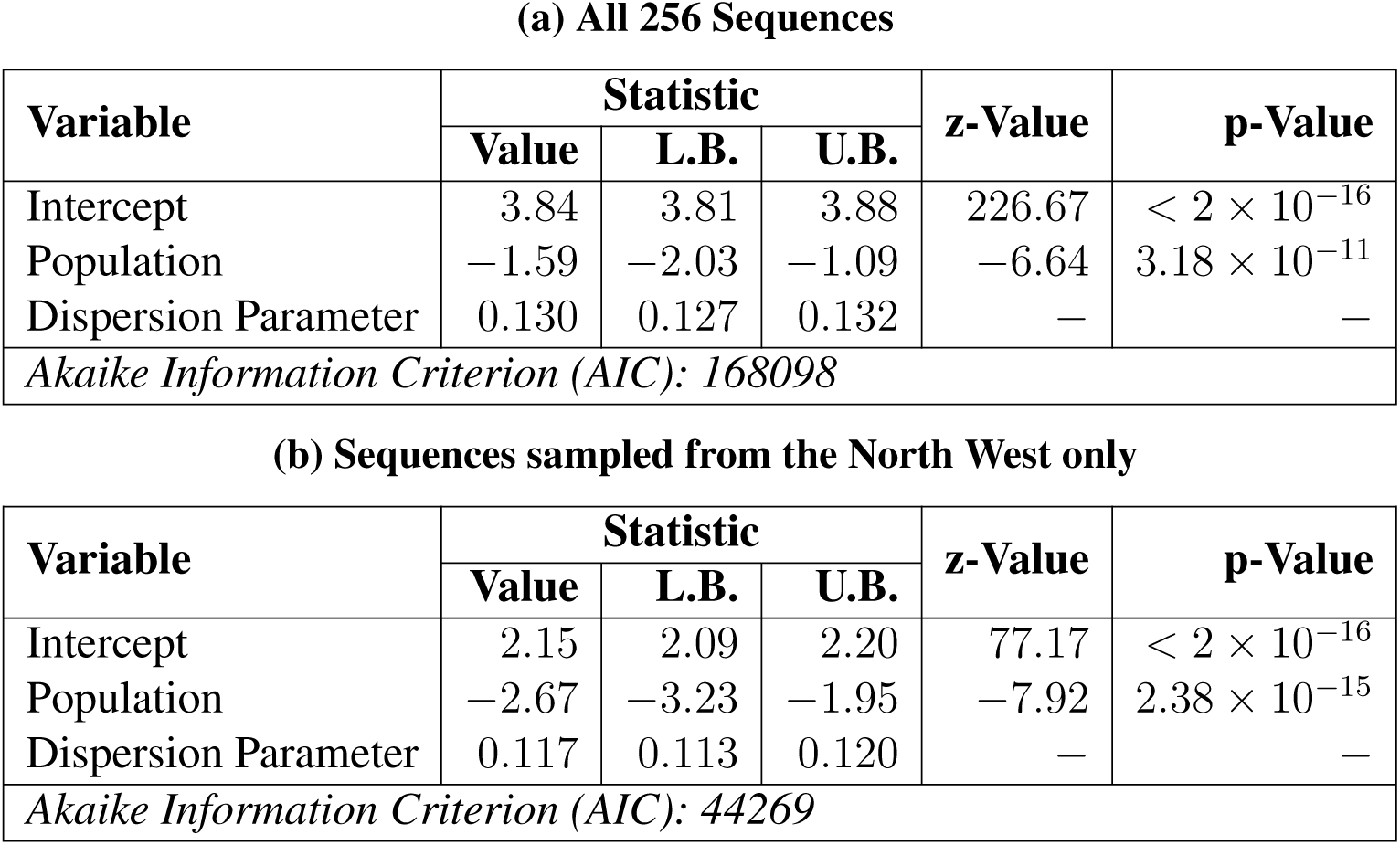
Results of Parametric Hypothesis Tests. Results of Pairwise Genetic Distance Parametric Hypothesis Tests using Negative Binomial Regression (see Section 3.5). Upper (U.B.) and lower (L.B.) bounds for each statistic produce a 95% confidence interval.

| Variable | Statistic |  |  | z-Value | p-Value |
| --- | --- | --- | --- | --- | --- |
|  | Value | L.B. | U.B. |  |  |
| Intercept | 3.84 | 3.81 | 3.88 | 226.67 | $< 2 \times 10^{-16}$ |
| Population | -1.59 | -2.03 | -1.09 | -6.64 | $3.18 \times 10^{-11}$ |
| Dispersion Parameter | 0.130 | 0.127 | 0.132 | — | — |
| *Akaike Information Criterion (AIC): 168098* |  |  |  |  |  |

**(b) Sequences sampled from the North West only**
| Variable | Statistic |  |  | z-Value | p-Value |
| --- | --- | --- | --- | --- | --- |
|  | Value | L.B. | U.B. |  |  |
| Intercept | 2.15 | 2.09 | 2.20 | 77.17 | $< 2 \times 10^{-16}$ |
| Population | -2.67 | -3.23 | -1.95 | -7.92 | $2.38 \times 10^{-15}$ |
| Dispersion Parameter | 0.117 | 0.113 | 0.120 | — | — |
| *Akaike Information Criterion (AIC): 44269* |  |  |  |  |  |

### 3.6 Molecular Analysis

To try to understand if adaptation was driving the main subclades in the epidemic clade (labelled A-F in 6), the mutations characterising them were analysed. These mutations are shown in Figure 9.

**Figure 9:**
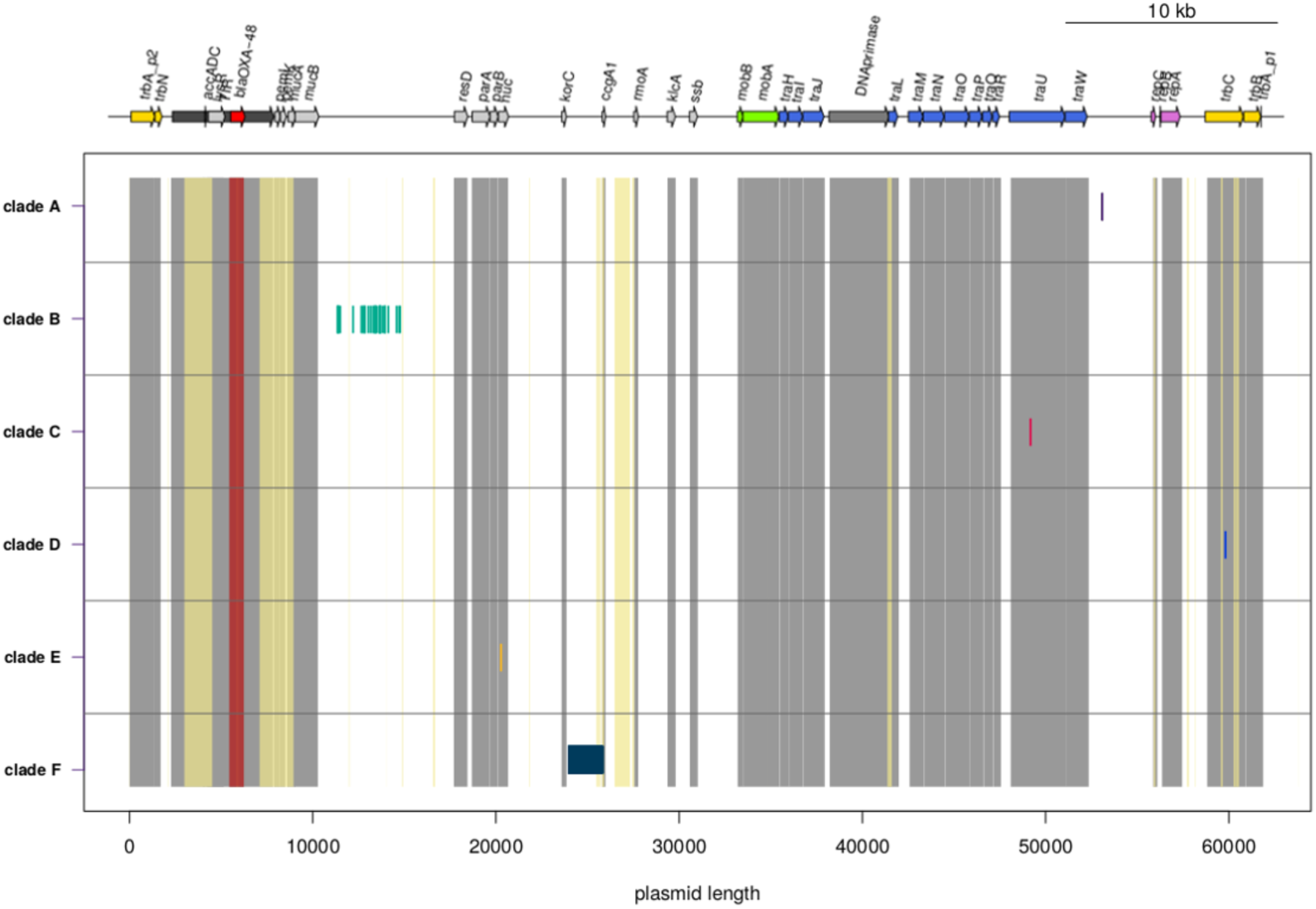
Mutations characterising subclades. Mutations characterising the subclades in Figure 6. The subclades have the same labels (A-F) and colours as in Figure 6. Known genes are labelled at the top of the figure, and the grey vertical bars indicate their presence in the plasmid. The yellow horizontal bars indicate that the region has been ‘masked’ (see `methods.rmd`). The region of the OXA-48 gene present in the plasmid sequence is shown with the red vertical bar. Subclade specific mutations are highlighted with short vertical bars in the same colour as the subclade highlights in Figure 6.

Four of the six subclades are characterised by just one nucleotide substitution (A, C, D, and E). The substitution characterising clade A is located in a non-annotated region. Substitutions characterising subclades C, D, and E are located in coding regions of the following genes: *DNA primase*, *traU* and *nuc*. At least two of these mutations are in genes that have been described to acquire mutations in previous studies: mutations in the *DNA primase* have been described also in de la Fuente et al 2022 [11], while mutations in the gene *nuc* have been described in Ledda et al 2021 [26].

Subclade B is characterised by 34 mutations in a 3500 bp non-coding region, from base 11356 to base 14754, most probably due to homologous recombination.

Subclade F is characterised by a 1567bp long deletion in a non-coding region (from base 24017 to base 25785). This deletion is overlapping with a hypervariable region in the same plasmid described in previous studies [26]. The fact that some of the mutations found in this study are in genes or intergenic regions already described as hosting mutations in other studies, suggests that those regions are more prone to mutations, and that these mutations might be adaptive.

## 4 Discussion

In this paper we have analysed a within-patient dataset of the pOXA-48 plasmid, finding that conjugation of identical plasmids in different bacterial hosts have a strong role in pOXA-48 dissemination. Reconstructing and comparing the plasmid in different sequences, we aimed at shedding light on the within-patient dynamics and evolution of this very well-conserved plasmid. We found that 85 of the 115 patients in our dataset carried at least two identical copies of the plasmid in different bacterial hosts. Only two of the 115 analysed patients had variants of the plasmid belonging to different clades of the tree, one from the “diverse” clade and one from the “epidemic” clade. Such a diversity in the plasmid sequence can only be explained by multiple infections.

This result has strong implications for public health investigations of pOXA-48 outbreaks: we find that in most patients the same allele of the plasmid is carried in different bacterial hosts. That means that, if we had sequenced just one sequence per patient, by solely focusing on the plasmid, we would have been able to identify at least 85/107 (79%) patients we found in the outbreak clade and which ones were carrying plasmids from the ‘diverse’ clade. On the other hand, if we had focused on sampling just resistant sequences from a specific bacterial host, like *Escherichia coli* or *Klebsiella pneumoniae*, we would have gotten a much more reduced vision of the outbreak, identifying 61 and 56 patients respectively in the epidemic clade. Other studies show that this is not true for other plasmids [13] which display a wider within-patient genomic variability. A better understanding of the within-patient plasmid variability and the role of transmission bottlenecks is clearly of paramount importance to develop effective investigation protocols for plasmid-mediated AMR outbreaks.

Of course, the dataset has some limitations: multiple sequences per patient were sequenced either if they carried different carbapenemase genes in the same bacterial host, or if they carried the same carbapenemase gene in a different bacterial host [16]. A very important part of the dynamics is therefore missing: how many times the same plasmid variant would have been identified in the same bacterial host had more than one colony pick per bacterial isolate been sequenced per patient. While knowing this would have changed the percentages in our results, it would not have changed the most important result of this analysis: that a patient very rarely acquires two different alleles of the pOXA-48 plasmid. We are, of course, not able to determine whether we find the same plasmid in two different bacterial host because the conjugation happened in the sampled patient or because the sampled patient was passed both hosts during the infection and the conjugation happened in another patient. This is, in a sense, not relevant as the main point to take home is that this plasmid is frequently conjugating and the odds of finding it in very different hosts are quite high, and this is what is important for devising appropriate screening protocols. Research on plasmid conjugation is at a turning point. One of the main problems so far is that many of the in vitro results cannot be even compared among themselves due to the different units in which they are expressed [18; 22]. New methods to study in vitro conjugation are being developed [18; 22] which will lead to a meaningful comparison of conjugation rate of different plasmids. Although in this case we haven’t been able to compute a rate for the within patient conjugation, we showed that it is widespread in our dataset. The limited variability of pOXA-48 has offered a unique possibility of avoiding all the sequencing and reconstruction issues affecting other plasmids to catch a glimpse of its within-patient dynamics [11]. With long read sequencing techniques becoming more and more reliable, such studies will be available for other plasmids and we will gain a clearer picture of within-patient plasmid dynamics.

This study shows how important it is to have a thorough understanding of the role of conjugation in plasmid-mediated AMR outbreaks [41]. It might inform sampling strategies in an outbreak investigation that can lead to a better understanding of the outbreak dynamics. By clarifying where the resistance is harboured and how often the plasmid conjugates, we could gain a better understanding of the effectiveness of potential approaches to mitigate an outbreak. Plasmid conjugation is the reason why plasmid-mediated AMR outbreaks are so concerning from the public health perspective. The more we understand about within-patient conjugation, the better tools we can develop for effective investigation of such outbreaks.

## Author Contributions

Conceptualization: A.L.; Data curation: K.L.H., A.L., F.G., T.N., L.L.; Formal analysis, Methodology, Investigation, Visualization: F.G., L.L., T.N., A.L.; Resources: K.L.H.; Funding acquisition: J.R., L.H., K.L.H.; Validation: D.W.; Writing: all authors.

## Abbreviations

AMR: antimicrobial resistance
OXA-48: gene of interest
pOXA-48: plasmid carrying the OXA-48 gene of interest
ST: sequence type
SNP: single nucleotide polymorphism
bp: base pair
kb: kilobase (pair)
CPE: carbapenemase-producing *Enterobacterales*
HGT: horizontal gene transfer

