## Supplementary Methods for "Plasmid conjugation drives within-patient plasmid diversity"

This document provides information on the code and analysis used.

---

#### Dataset

##### Overview

Data was provided by Katie L Hopkins (Antimicrobial Resistance and Healthcare Associated Infections (AMRHAI) Reference Unit - Specialised Microbiology and Laboratories Division - UK Health Security Agency and Antimicrobial Resistance and Prescribing - HCAI, Fungal, AMR, AMU & Sepsis Division - UK Health Security Agency).

The dataset consists of 527 sequences. These come from 243 patients. Each patient has between 2 to 5 sequences. The sequences for each patient were either:

1) sequences with the same resistance that were found in different bacterial hosts in the same patient, or
2) sequences with the same bacterial hosts but with different resistances found in the same patient.

##### Filtering the dataset

We focus on patients with sequences with the *OXA-48* gene. Sequences were lab tested for containing OXA-48 (and other genes encoding resistance mechanisms to carbapenems, such as NDM and VIM).

We filter down to patients with at least one sequence determined to contain OXA-48. This leaves us with 121 patients with 269 sequences between them.

Later on, we will further remove sequences due to later issues with assembly, alignment, and data quality (see "Deduplication" and "Checking consistency" sections). This will eventually leave us with 115 patients with 256 sequences between them.

##### Sequence Isolation

The `fastq` files (containing the short read sequence data) contain duplicates. Some of the files were duplicates due to either: - the *same* sample was sequenced twice - (we would have kept these) - or the sequenced file did not correspond to the lab assigned species (in this case nobody knows what was sequenced).

It was safer to exclude duplicated samples in the analysis (see Deduplication section).

### Sequence reconstruction

To be sure that we had the best plasmid reconstruction possible, we reconstructed the plasmid sequences using two parallel approaches: 1) "hybrid de-novo" approach, and 2) a standard "on template"/"reference-based" reconstruction.

For both approaches, the plasmid sequence we used as a template has refseq accession number `JN626286.1`, published by Poirel,L., Bonnin,R.A. and Nordmann,P in 2011. - This can be accessed in the NCBI Nucleotide library.

#### Method 1: Hybrid de-novo Reconstruction

##### Step 1: SPAdes

- We run the `Plasmid SPAdes` program with standard options, using the High Performance Computing Cluster (HPC).
- We pass as inputs all of the `fastq` files for each sample in the dataset
- The link to the SPAdes github repository is: https://github.com/ablab/spades.
- The bash job file to run this has been included in the cell below:

Language: bash

```bash
#!/bin/bash
#$ -N SPAdes
#$ -S ./logsAnderrors/
#$ -pe multithread 8
#$ -l h_vmem=8G

source $HOME/.bash_profile
source /etc/profile.d/modules.sh
module purge
module load python/2.7
module load spades

spades.py -1 input1 -2 input2 -o output \
--plasmid \
--phred-offset 33 \
--careful \
--cov-cutoff auto \
--threads 8 \
--memory 8 \
-k '21,33,55,77'
```

- The outputs were stored in a folder called `SPAdesAss`, and the `SPAdes` output is saved in a unique folder for each sample.
- The `fasta` output files we are interested in are saved as `contigs.fasta` within each sample's folder.
- We collate all of these `contigs.fasta` files into a new folder and rename them after their sample `molisId` using the python script below:

Language: python

```python
import os, shutil, logging

input_folder = r"~/PlasmidSpadesAssCO"
```

```
output_folder = r"~/all_contigs_files"
log_filepath = r"~/all_contigs_files/rename_contigs.log"
contig_folder_prefix = "COPlasmidSpadesAss"

# Setup logging
logging.basicConfig(filename=log_filepath, level=logging.INFO)

# Get names of contig folders
contig_folders = [x for x in os.listdir(input_folder) if x.startswith(contig_folder_prefix)]

# Iterate over contig folders and copy to new destination with new name
for c in contig_folders[:5]:
    new_filename = f"{c.replace(contig_folder_prefix, '')}.fasta"
    destination_filepath = f"{output_folder}/{new_filename}"
    source_filepath = f"{input_folder}/{c}/contigs.fasta"
    if not os.path.exists(destination_filepath):
        try:
            shutil.copy(source_filepath, destination_filepath) # Copy files
            # os.rename(source_filepath, destination_filepath) # Rename files
            logging.info(f"Successfully copied '{source_filepath}' to '{destination_filepath}'")
        except:
            logging.info(f"Unable to copy '{source_filepath}' to '{destination_filepath}'")
    else:
        logging.info((
          f"File '{new_filename}' already exists." +
          f"Not attempting to copy '{source_filepath}' to '{destination_filepath}'"
        ))
```

##### Step 2: Creating dotplots

- We tried using both `lastz` and `mummer4` to assemble/align from the SPAdes outputs.
- These programs output coordinate files that we can plot dotplots from.
- We decided not to use lastz, and use mummer4 instead.
- The link to the mummer4 homepage is https://mummer4.github.io/.
- The steps to run the mummer4 pipeline are:

  1. Run `nucmer`
  2. Run `delta-filter` on outputs of 1.
  3. Run `show-coords` on outputs of 2.
  4. (Optional) Run `show-aligns` also on outputs of 2. (This isn't exactly needed but is nice to have)
  5. Run python script to plot dotplots from output of 3.

- Note: `nucmer`, `delta-filter`, `show-coords`, and `show-aligns` are command line tools that come with mummer4.
- The bash job file for running the mummer4 pipeline (steps 1-4) is included below:

Language: bash

```
#!/bin/bash
#SBATCH --job-name=mummer_pipeline
#SBATCH --mem=12GB
#SBATCH --time=120:00:00

### SETUP ###
# Home directory:
home="~"
```

```bash
# Plasmids directory:
plasmids_path="~"

# Query Contigs directory
query_contigs_path=$plasmids_path/all_contigs_files
# Reference Contig to compare to
ref_contig_path=$plasmids_path/pOXA48.fasta

# Output directories:
output_path=$query_contigs_path/mummer_outputs
delta_path=$output_path/delta_files
filtered_delta_path=$output_path/filtered_delta_files
coord_path=$output_path/filtered_coordinate_files
aln_path=$output_path/filtered_alignment_files

# Create output directories (if not exists)
mkdir -p $output_path
mkdir -p $delta_path
mkdir -p $filtered_delta_path
mkdir -p $coord_path
mkdir -p $aln_path

# Select conda environment:
source ~/anaconda3/etc/profile.d/conda.sh

ref_contig=`basename "$ref_contig_path" .fasta`;

# Shell option to expand unmatched shell glob to expand to nothing
# i.e. if there are no files to iterate over, don't perform `do` action
shopt -s nullglob

unset IFS # Reset default delimiter for arrays

echo "Reference contig file: ${ref_contig_path}"

# Iterating over contig files:
for FILE in "$query_contigs_path"/*.fasta; do
  echo "-----"
  echo "Query contig file: ${FILE}"
  query_contig=`basename "$FILE" .fasta`;
  echo "Applying nucmer"
  nucmer --delta="${delta_path}/${ref_contig}_${query_contig}.delta" "$ref_contig_path" "$FILE"
  echo "Applying delta-filter"
  delta-filter \
    -q "${delta_path}/${ref_contig}_${query_contig}.delta" >
    "${filtered_delta_path}/${ref_contig}_${query_contig}.filtered.delta"
  echo "Applying show-coords to filtered"
  show-coords \
    -T \
    -rcl \
    "${filtered_delta_path}/${ref_contig}_${query_contig}.filtered.delta" >
    "${coord_path}/${ref_contig}_${query_contig}.filtered.coord"
  echo "Applying show-aligns to filtered"
  IFS=$' ' # Change delimiter to space
  LINES=`grep ">" "${filtered_delta_path}/${ref_contig}_${query_contig}.filtered.delta" |
    awk '{sub("^>","");print}'`
  IFS=$'\n' # Change delimiter to newline
  for X in ${LINES}; do
```

```
    IFS=$' ' read -r -a array <<< "$X"
    rId=${array[0]} # reference node id
    qId=${array[1]} # query node id
    echo "Comparing ${rId} to ${qId}"
    show-aligns \
      -r "${filtered_delta_path}/${ref_contig}_${query_contig}.filtered.delta" "${rId}" "${qId}" >>
      "${aln_path}/${ref_contig}_${query_contig}.filtered.${qId}.aligns";
  done;
done;
```

To create the dotplots (step 5 above), use the following python script:

Language: python

```python
import os, re
import pandas as pd
from matplotlib.axes import Axes
import matplotlib.pyplot as plt
import numpy as np

# directory of show-coords outputs to create dotplot plots from
# - ${coord_path} in the bash script above
path_to_show_coords_outputs = ""

# directory to save dotplot plots to
save_directory = ""

reference_contig_name = "pOXA48"
show_coords_filenames = [x for x in os.listdir(path_to_show_coords_outputs) if x.endswith(".coord")]

def plot_lines(
    ax: Axes,
    x0: list[float],
    x1: list[float],
    y0: list[float],
    y1: list[float],
    plot_args: dict = {"linewidth": 2, "markersize": 3}
    ) -> Axes:
    """
    Returns plot of lines (x_0, y_0) to (x_0, y_0) for x_0, x_1, y_0, y_1 in zip(x0, x1, y0, y1).

    Instead of individually plotting lines, we plot one line object broken up by NaN values.

    E.g. say we have the following
        - ``x0 = [0, 0, 10]``
        - ``x1 = [5, 1, 25]``
        - ``y0 = [9, 8, 21]``
        - ``y1 = [14, 9, 36]``

    This function will plot on the `ax` Axes three lines:
        - (0,9) to (5,14)
        - (0,8) to (1,9)
        - (10,21) to (25,36)

    This is done by plotting the following:
        ``plt.plot(x = [0,5,NaN,0,1,NaN,10,25,NaN], y = [9,14,NaN,8,9,NaN,21,36,NaN])``

    Parameters
```

```
    ----------
    ax : matplotlib.axes.Axes
        Axes object to add artists to
    x0, x1, y0, y1 : list of float
        List of coordinates of lines (x0, y0) to (x1, y1)
    plot_args : str, optional
        kwargs to pass to plot function

    Returns
    -------
    matplotlib.axes.Axes
        Dotplot plot
    """

    # Create list of x and y values separated by NaNs
    x = np.array([[start, end, np.nan] for start, end in zip(x0, x1)]).ravel()
    y = np.array([[start, end, np.nan] for start, end in zip(y0, y1)]).ravel()

    # Plot x, y onto axes:
    ax.plot(x, y, "o-", **plot_args)

    return ax

# iterate over all files in show_coords_filenames
for filename in show_coords_filenames:
    # When we turn to class, all this should be moved to relevant dotplotter...
    df = pd.read_csv(
        os.path.join(path_to_show_coords_outputs, filename),
        sep="\t", # Tab delimited file
        skiprows=[0,1,2,3], # Ignore the first 4 lines of file
        index_col=False, # No index column contained in file
        usecols=[0,1,2,3], # Only read the first 4 columns
        names=["x0", "x1", "y0", "y1"] # Name of columns
    ).assign(hue=0)

    sample_name = ".".join(filename.split(".")[:-1])

    # Create Plot
    fig, ax = plt.subplots(figsize=(7, 7))

    # Plot lines by each group
    for _, d in df.groupby("hue"):
        ax = plot_lines(ax, d.x0, d.x1, d.y0, d.y1)

    # Additional plot features
    ax.axis("square") # equal axes and square plot
    ax.set_xlabel(reference_contig_name)
    ax.set_ylabel(sample_name)

    # Save plot:
    plt.savefig(os.path.join(save_directory, filename.replace(".coord", ".png")))
    plt.close()
```

##### Step 3: Manually assessing dotplots

- We manually checked through all of the dotplots.
- Our rule of thumb for a "good" sample was:

1. coverage throughout, with some freedom either end (i.e. for `JN626286.1`, this has ~60k base pairs, so we'd want coverage from ~4k to ~55k. This means diagonal lines must lie over this range on the x axis).
2. diagonal lines must touch from x axis (with some freedom - e.g. either end, and longer diagonal lines may be slightly broken - only allow for very small gaps)
3. coverage between 5k-10k is critical - this is where `OXA-48` is present in `JN626286.1`

- This will help us to get a good idea of whether our reference based assembly (second reconstruction method) is valid or not.

##### Step 4: Checking `contigs.fasta` with `mlst`

- **See: https://github.com/tseemann/mlst**
- Once samples were deduplicated and potentially contaminated samples were excluded, the results from MLST agreed with the species assignation we were provided. SO we kept the species identification that we were provided by K. Hopkins.
- there is a file with all the species identification andthe MLST results. *add the file name here*
- NOTE: I do not think this step belongs here, it belongs out of the hybrid assembly pipeline, but it is not a big deal.

#### Method 2: Reference-based assembly

##### Step 1: Indexing with BowTie2

We run BowTie2 on the HPC to index the OXA-48 reference plasmid.

Language: bash

```bash
#!/bin/bash


source $HOME/.bash_profile
source /etc/profile.d/modules.sh
module purge
module load bowtie2

bowtie2-build -f pOXA48.fasta OXA48
```

##### Step 2: Alignment with BowTie2

Then we aligned the sequence reads on the reference using Bowtie2 on each set of `fastq` files for each sample.

Language: bash

```bash
#!/bin/bash
#$ -N bowtie2
#$ -S ./logsAnderrors/
#$ -pe multithread 8
#$ -l h_vmem=8G


source $HOME/.bash_profile
source /etc/profile.d/modules.sh

module purge
```

```
module load bowtie2

bowtie2 -x OXA48 -q -1 input_1 -2 input_2 -S outputFile
```

##### Step 3: Build consensus using Samtools

Then we run samtools consensus in two different ways. One with the default options and one with cutoffs on the base quality and alignment quality.

The default one was as in the following script

Language: bash

```bash
#!/bin/bash
#SBATCH --job-name=samtoolsConsensus.sh
#SBATCH --mem=12GB
#SBATCH --time=120:00:00

# Running simple samtools script to get the consensus sequence
for FILE in `ls Bowtie2/*.sam |awk '{gsub(/.sam/,"");print}'`; do
  samtools consensus -f fasta -a -o "$FILE".pl.fst "$FILE".pl.bam ;
done ;
```

- Its output is called `MolisID.pl.fst`.
- The concatenated file using all the files obtained using this scripts + the reference is called all.fst
- The `samtools` consensus script with stronger parameters is here

Language: bash

```bash
#!/bin/bash
#SBATCH --job-name=testrun-samtools.sh
#SBATCH --mem=12GB
#SBATCH --time=120:00:00
#SBATCH --partition=PARTITION-NAME

# Running simple samtools script to get the consensus sequence
for FILE in `ls Bowtie2/*.sam |awk '{gsub(/.sam/,"");print}'`; do
  samtools consensus -f fasta -a \
  --min-MQ 30 \
  --min-BQ 30 \
  --show-del yes \
  -o "$FILE".ID.fst "$FILE".pl.bam ;
done ;
```

- We introduced a minimum quality for base quality and alignment quality of 30.
- We also asked it to show deletions, but this option could be relaxed.
- The output is called `MolisID.ID.fst`.
- The concatenation of all the files and the reference is called `all_ID.fst`.

##### Step 4: Cleaning/editing consensus

- We then started cleaning the concatenated file to be aligned using R.

Language: r

```r
library(tidyverse)
library(ape)


# all.fst the file reconstructed using samtoools consensus with standard options

seq <- read.dna("all.fst", format = "fasta")
seqc <- as.character(seq)

numn <- t(sapply(seqc, function(x) {
    mean(x == "n")
}))

hist(numn)
hist(numn, breaks = 100)

seqc_clean <- lapply(seqc[which(numn < 0.2)], function(x) {
    x[x != "n"]
})
# write.dna(as.DNAbin(seqc_clean),file='all_clean.fst',format='fasta',colsep='',blocksep='')

seqc_clean_withN <- seqc[which(numn < 0.2)]
# write.dna(as.DNAbin(seqc_clean_withN),
# file='all_clean_withN.fst',format='fasta',colsep='',blocksep='')
```

This leaves us with two files to align: - one where we removed all the `N`s - one where we kept all the `N`s that samtools conssensus put there.

Having the `N`s might be important because it means that there was a base, just not a strong enough signal as to determine which base it is.

We also run the same script using the quality filtered consensus files.

### Deduplication

Of the 310 `fastq` files, 46 were duplicates (pairs of data from the *same* plasmid sequence samples ending in a '.0' or '.1').

In case of a duplicated file, we used the following algorithm to choose one of the two available samples:

1. **species**: if one of the samples was classified by MLST as the same species as the one K. Hopkins gave us and one did not -> the sample of a different species was discarded irrespective of plasmid presence/absence
2. **plasmid presence**: if both samples were classified by MLST as the SAME species K. Hopkins assigned them to AND one of them had the OXA-48 plasmid and one did not have the plasmid, then we kept the plasmid-carrying sample
3. **plasmid length**: if both plasmids belonged to the same species AND both had the OXA-48 plasmid, then the shorter reconstructed plasmid was discarded (often the difference was 10s to 100s basepairs, so probably there was not much of a difference, but we had to enforce a general rule)

PATIENT199 had 2 couples of duplicated samples. In the first couple of duplicated samples (PAT199-S_431.0 and PAT199-S431.1), neither of them were assigned to the same SPECIES assigned by K. Hopkins, so both were discarded. Discarding them would have left us with just one sample (which was again a duplicated sample), so we choose to discard the entire patient.

The list of 48 samples that were excluded in the deduplication process is the following:

```
"PAT09-S_022.1"
"PAT111-S_243.0"
"PAT111-S_244.0"
"PAT120-S_264.1"
"PAT125-S_275.0"
"PAT125-S_276.1"
"PAT128-S_282.1"
"PAT129-S_284.1"
"PAT155-S_336.0"
"PAT156-S_338.1"
"PAT157-S_340.0"
"PAT157-S_342.0"
"PAT161-S_350.1"
"PAT174-S_377.0"
"PAT174-S_376.1"
"PAT183-S_397.0"
"PAT199-S_430.0"
"PAT199-S_431.0"
"PAT199-S_430.1"
"PAT199-S_431.1"
"PAT201-S_436.1"
"PAT21-S_050.0"
"PAT21-S_051.0"
"PAT211-S_457.0"
"PAT211-S_458.1"
"PAT214-S_464.1"
"PAT222-S_481.1"
"PAT23-S_054.1"
"PAT230-S_497.1"
"PAT235-S_509.0"
"PAT240-S_520.1"
"PAT28-S_064.0"
"PAT33-S_074.0"
"PAT37-S_083.1"
"PAT37-S_084.1"
"PAT37-S_085.1"
"PAT45-S_102.1"
"PAT49-S_110.1"
"PAT51-S_114.1"
"PAT52-S_117.1"
"PAT56-S_124.0"
"PAT56-S_125.1"
"PAT56-S_126.1"
"PAT72-S_159.1"
"PAT80-S_176.1"
"PAT90-S_197.0"
"PAT93-S_206.1"
"PAT96-S_212.1"
```

We excluded a total of 48 samples, 4 of which were from PATIENT199 alone. We keep the other non-discarded sample and remove the '.0' or '.1' suffix. (e.g. we discard PAT93-S_206.1 but keep PAT93-S_206.0 and rename this sample PAT93-S_206) After deduplication, we were left with 262 samples (and fastq files) from 118 patients. The script used to deduplicate all the files is `scripts/Deduplicate.R`

### Files

The new files for reprocessing many of the analysis above after deduplication are as follows:

- epi data that came with the files:
  - Original file `20210615-dataKatieMolis.csv`.
  - **Deduplicated file `finalEpiDataRef.csv`** (relevant data from this has been merged into `supplementary_data.csv`)
- spreadsheet with assembly data:
  - Original file `PlasmidsToDelete-AssemblytoClean.csv`.
  - **Deduplicated file `deduplicatedAssembly.csv`**
  - **Cleaned with MOLIS IDs removed: `supplementary_data.csv`**
- full alignment of all the reconstructed plasmids.
  - Original file "all_clean_id_aligned.fst".
  - **Deduplicated file `deduplicated_id_seq_clean_aligned.fasta`**
  - **Cleaned with MOLIS IDs removed: `aligned_sequences.fasta`**
  - Aligned file (see below) `REalign_deduplicated_id_seq_clean.fst`.
  - File with Patient Ids instead of MOLIS IDs `REalign_deduplicated_id_seq_clean_PAT.fst`
- Alignment above with problematic regions masked off (See below for details):
  - **Deduplicated file `Final_alignment_masked_PAT_50bases_0.1samples.fst`**
  - *NOTE: the deduplicated file has had the MOLIS_ID subsituted with the PAT-Sample_IDs*

### Scripts

R scripts that run the analysis on deduplicated files:

- script for making all the plots of epidemiological features of dataset (Figure 1 in paper):
  - `plotting_epidemiological_features.R`
- script for making plot of genetic distances for between and within-patient (Figure 4 in paper):
  - `plotting_pairwise_genetic_distance_distributions.R`
- script for tree figures:
  - `plotting_trees.R`
- script for redoing the tree (see forward for details):
  - `scripts/Re-alignment4betterTree.R`
  - `scripts/mask_mutations_mainclade_OXA48.R`
- script for within patient genetic distance analysis:
  - `scripts/withinPatDistNewDedu.R`
- script for the genomic analysis:
  - `scripts/Genomic_analysis.R`

- script for parametric hypothesis tests:
  - `scripts/negative_binomial_glm_hypothesis_test.R`
- script for non-parametric hypothesis tests:
  - `scripts/non_parametric_hypothesis_test.py`
- script for preparing the files for `Genomic_analysis.R`:
  - `scripts/AugurJsonScript.R`

### Checking consistency

Of the 262 samples:

- 2 have nothing at all in any pipeline (PAT07-S_014 and PAT73-S_162)
- 1 further sample was lost by the PS-M pipeline only (PAT241-S_521)

The Plasmids-SPAdes-mummer pipeline yielded 260 dotplots, 17 of which were empty.

The samtools pipeline produced an alignment containing 262 (+ the reference genome) aligned sequences (of length at least 49,806 bps out of 61,881 bps of the reference plasmid).
The 28 samples for which we were not able to get a reconstructed sequence were also mostly lacking a proper reconstruction in the other pipelines.

We do encounter 2 exceptions, PAT202-S_437 and PAT202-S_438, which seem to be having insertions in the last part of the plasmid, but also maybe experiencing:

- poor sequencing
- too many contigs
- large gap(s) (in particular in the middle of the sequence)

We remove the samples from patient PAT202.

3 samples seem to have the OXA-48 gene and nothing else:

- PAT09-S_020 has the OXA-181 allele, which is usually located in the chromosome.
- PAT182-S_394 is an E.coli ST38 sample, where the OXA-48 gene is located in the chromosome. We remove this sample. Patient PAT182's other sample (S_395) is not a sample witn OXA-48, so we remove Patient PAT182 entirely.
- PAT92-S_204 is a Citrobacter sample.

All of this is in a file called ***`deduplicatedAssembly.csv`***. The cleaned version of this file (given) is ***`supplementary_data.csv`***.

The R pipeline to obtain this file is in `checkSamples.R`.

- We visually checked the dotplots obtained with mummer4 to be sure that those samples had problems.
- We also checked the dotplots of the samples in the longer 'Epidemic' clade.

A further 3 patients are removed from the dataset: PAT186, PAT230, PAT243. The samples for these 3 patients either: - have a partial plasmid sequence, so we removed the sample, or - had two samples taken on the same day and were very similar but in different hosts (or the host assignation was not univocal) and we had no way of choosing a "trusted" representative, so we flushed both of them.

This leaves us with 256 samples from 115 patients.

### Alignment and Tree building

#### Step 1: alignment

We aligned all the assembled files using mafft with default options.

Language: bash

```bash
mafft deduplicated_id_seq_clean_aligned.fasta > REalign_deduplicated_id_seq_clean.fasta
```

#### Step 2: cleaning the alignment

The alignment has several regions where the sequences did not align properly. Mutations closely adjacent to gapped regions cannot be trusted as could be resulting from misalignments in the gapped region. To mask these regions we used an algorithm in two steps:

1. IF in a certain position gaps accounted for more than 10% of the sequences (n=24) we masked all the mutations in all samples within 50 bases from this position
2. IF in a certain position gaps accounted for less than 10% of the sequences (n<24) we masked we masked all the mutations within 50 bases from this position but only in the samples where the gaps were present.

The rationale for this choice was that if gaps were present in more than 10% of the sequences we were likely dealing with a variable region that could not be properly aligned everywhere, if the gaps were present in fewer than 10% of the sequences we asssume that the problem was lying only in the affected sequences and not intrinsically in the genomic region.

In this way we ended up masking 5989 sites out of 63797 of the entire alignment (9.3% of the sites)

The script to do this is *maskingAlignment0.2.R*

The output is *Final_alignment_masked_PAT_50bases_0.1samples.fst*

All the masked intervals are saved in a table called *table_masked.csv*

#### Step 3: making the tree

We built the tree using IQtree under the GTR substitution model (cite Tavare 1986) with the discrete Gamma model (cite Yang, 1994) with the 4 default rate categories to correct for rate heterogeneity across sites .

Since the tree is a binary object and in this dataset there are many identical sequences, we did not want the tree to show small branches where there were none. For this reason we used the flag `-blim` to set the length of such branches very short, invisible to the plotted tree.

Language: bash

```bash
iqtree -s Final_alignment_masked_PAT_50bases_0.1samples.fst -m GTR+G -blmin 0.00000001
```

### Assigning bacterial host

Bacterial hosts were assigned via "maldi tof" in the lab, and then confirmed and detailed "in silico". The species characterization resulting from this process is here. It covers all the 310 samples.

Language: r

```r
library(tidyverse)
library(ape)

sttab_OXA48 <- as_tibble(read.csv("sttab_OXA48.csv"))

sttab_OXA48 %>%
    group_by(SP) %>%
    count() %>%
    arrange(desc(n))
```

We decided to check the bacterial host species by running `MLST` on the `SPAdes` assembly (*not* described above). `MLST` produced results for 305 of the 310 samples and the species assigned were quite varied:

Language: r

```r
library(tidyverse)
library(ape)

mlst_output_filepath <- "mlst_output_Fan_clean.csv"

newMLST <- as_tibble(read.csv(mlst_output_filepath)) %>%
    mutate(SP = pubmlst_scheme_name) %>%
    mutate(SP = ifelse(SP == "kpneumoniae", "Klebsiella_pneumoniae", SP)) %>%
    mutate(SP = ifelse(SP == "koxytoca", "Klebsiella_oxytoca", SP)) %>%
    mutate(SP = ifelse(SP == "ecloacae", "Enterobacter_cloacae", SP)) %>%
    mutate(SP = ifelse(SP == "ecoli", "Escherichia_coli", SP)) %>%
    mutate(SP = ifelse(SP == "paeruginosa", "Pseudomonas_aeruginosa", SP)) %>%
    mutate(SP = ifelse(SP == "senterica", "Salmonella_enterica", SP)) %>%
    mutate(SP = ifelse(SP == "cfreundii", "Citrobacter_freundii", SP))

newMLST %>%
    group_by(SP) %>%
    count() %>%
    arrange(desc(n))
```

### Within-patient analysis

The R scripts for the within patient analysis are in **"withinPatDistNewDedu.R"** in the repository.

The statistical analysis of the distribution of SNPs comparing between patient and within patient population has been done in the following notebooks in the repository (in the `scripts` folder):

- `negative_binomial_glm_hypothesis_test`
- `non_parametric_hypothesis_test.py`

The data used is created in the `plotting_pairwise_genetic_distance_distributions.R` script, and is saved as `genetic_distance_table.csv`.

### Genomic analysis

We wanted to understand which mutations characterised each subclade. First off we need to add node labels to the tree, as we will need it later on.

Language: r

```
library(tidyverse)
library(ape)

masked_tree_Pat <- read.tree("Final_alignment_masked_PAT_50bases_0.1samples.fst.treefile")
masked_tree_Pat$node.label <- paste("node", 1:masked_tree_Pat$Nnode, sep = "-")
write.tree(masked_tree_Pat, "Final_NODES_alignment_masked_PAT_50bases_0.1samples.fst.treefile")
```

Then we run `augur ancestral` using the following script:

Language: bash

```
augur ancestral \
--tree Final_NODES_alignment_masked_PAT_50bases_0.1samples.fst.treefile \
--alignment Final_alignment_masked_PAT_50bases_0.1samples.fst \
--root-sequence pOXA48.fst \
--keep-ambiguous \
--output-node-data Final_NODES_augur.output_dedu.data.json \
--output-sequences Final_NODES_augur.output_dedu.seq.fst
```

The results from Augur ancestral were processed using the scripts `AugurJsonScript.R` and `Genomic_analysis.R` to produce figure 9
